# Hippocampal Representational Drift Persists in a Stable Multisensory Virtual Environment

**DOI:** 10.1101/2025.04.14.648740

**Authors:** Jason R. Climer, Heydar Davoudi, Jun Young Oh, Daniel A. Dombeck

## Abstract

Experiments tracking hippocampal place cells in mice navigating the same real environment have found significant changes in neural representations over days. However, there is currently a debate over whether such “representational drift” serves an intrinsic function, such as distinguishing similar experiences occurring at different times, or is instead observed due to subtle differences in the sensory environment or behavior. Here, we used the experimental control offered by a multisensory virtual reality (VR) system to determine that differences in sensory environment or behavior do not detectably change drift rate. We also found that the excitability of individual place cells was most predictive of their representational drift over subsequent days, with more excitable cells exhibiting less drift. These findings establish that representational drift occurs in mice even with highly reproducible environments and behavior and highlight neuronal excitability as a key factor of long-term representational stability.

## Introduction

Place cell ensembles in the hippocampus had long been thought of as the stable substrate for spatial long-term memories^1^. Yet, experiments tracking place cells in mice navigating the same environment over days challenged this idea with evidence of significant changes in the representations, a phenomenon recently termed “representational drift”^2–7^. Hippocampal representational drift could serve as a mechanism to distinguish similar experiences occurring at different times^8,9^ or provide computational benefits such as continual learning^8^, generalization^10,11^, allowing for redundancy^12^ or be a side effect of computations that allow for these benefits^10,12^. Conversely, drift could simply be a read-out of subtle differences in behavior, such as running speed^13–16^, or the sensory environment, such as olfactory cues^17–19^, that are difficult to measure and control yet highly notable and perceptible to the animal. This latter hypothesis that drift occurs due to subtle sensory or behavioral variabilities has gained some support from recent observations^13^ and analyses^20^. For example, in the hippocampus CA1 of Egyptian fruit bats, little representational drift was observed over days when the behavior (i.e., flight path) was highly stereotyped^21^. Importantly, however, few experiments controlling for subtle behavioral and environmental variabilities have been performed, making it unclear whether such variabilities could provide a general explanation for representational drift. Therefore, we used the precise sensory control afforded by a multisensory (visual+olfactory) VR system^22,23^ and the behavioral control achieved by head-fixed mice running through the environment using a linear treadmill to determine if subtle behavioral and sensory variabilities impact the rate of representational drift in hippocampal CA1 of mice.^24^

## Results

### Hippocampal representational drift occurs in mice navigating a familiar virtual environment

Representational drift in mouse CA1 place cells has been observed in real^2,3,6,7,9,25,26^ and virtual^4,5,27^ environments with different environment sizes and shapes, treadmill types (e.g., spherical with motion constrained to one dimension^4^ or cylindrical^27^), sensory cues, and reward conditions. Therefore, we first sought to confirm that hippocampal CA1 drift occurs in our apparatus, consisting of a virtual linear track and a one-dimensional cylindrical treadmill (Fig. 1a-b)^28–31^, and to quantify the drift rate with many previously used metrics^3–5,25,26^.

**Figure 1:**
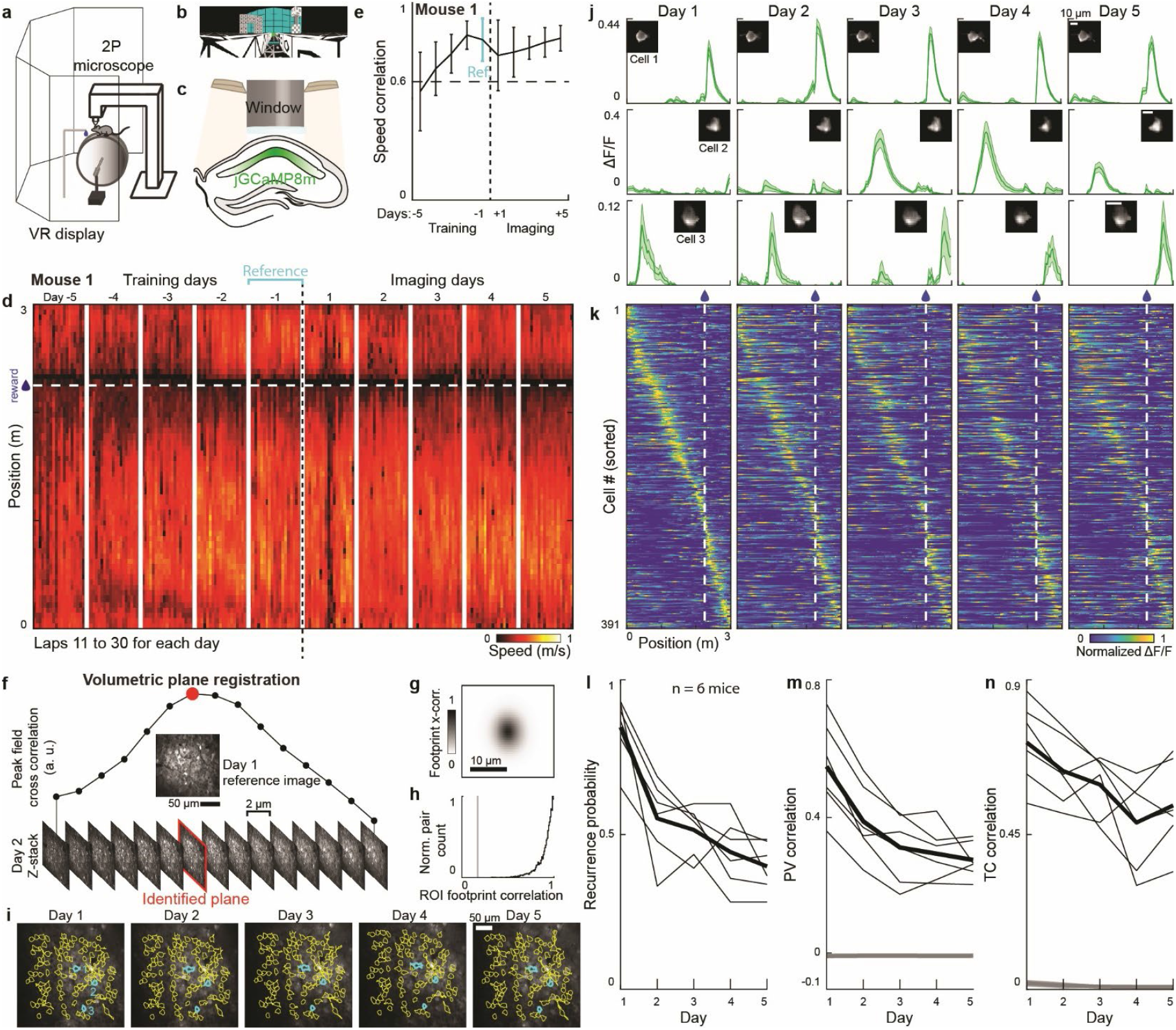
Hippocampal representational drift occurs in a familiar virtual environment. a) The experiment setup: head-fixed mouse running on a cylindrical treadmill in front of a five-panel VR display and receiving water rewards during 2P imaging. b) The mouse navigates a 3-m one-dimensional virtual linear track. c) 2P imaging of dorsal CA1 neurons using GCaMP8m through a hippocampal window. d) The speed profile of a trained exemplar mouse performing the task in a familiar environment over five training and five imaging days. Each column shows a single-track traversal lap. The horizontal dash line and blue droplet indicate the water reward location, and the vertical white lines separate each day. e) The speed correlation of laps 11-30 on each day to the mean of laps 11-30 on the reference day (cyan). A horizontal dashed line indicates the correlation threshold. f-i) Across-day volumetric plane registration method. f) A series of z-stacks (15 slices, 2 µm apart) were obtained around the Day 1 imaging plane. Maximum cross-correlation values for the 15 z-planes to Day 1 imaging plane were calculated and the peak selected as the matching plane (red dot). g) Cross-correlation of spatial footprints for all ROI pairs aligned across days. h) Histogram of spatial footprint correlations for all ROI pairs aligned across days. Cross-cell shuffles are shown in gray. i) The selected imaging planes in an example mouse across five days. Cell ROIs were identified using Suite2p^45^ and aligned across days using CellReg^46^. Three example ROIs identified on all five days highlighted in cyan. j) Three example place cells from (i) showing heterogeneous place tuning curves across days. Mean DF/F versus track position is shown for each cell across days. The first place cell is stable over days, the second place cell field emerges on day 3, and the third place cell field shifts location across days. The morphologies of these three cells are shown in insets, confidence intervals show lap-by-lap standard error of the mean (SEM). k) Sorted and cross-validated population heatmaps of 391 cells (6 mice) identified as place cells on Day 1. Each horizontal row shows the mean DF/F of each neuron vs track position. Day 1: the rows are sorted based on the place field locations on odd laps, and the even lap mean DF/F is plotted. Days 2-5: same 391 cells sorted according to day 1 order, plotting mean DF/F for even laps on each indicated day. l-n) Population representational drift measured by recurrence probability (l), population vector (PV) correlation (m), and tuning curve (TC) correlation (n). Thin line, each mouse; thick line, mean across mice; grey line, random shuffle.

To reduce behavioral variables outside the recording session that may affect hippocampal drift^32,33^, we singly housed mice in identical, unenriched cages (e.g., no exercise wheels), transported them in enclosed dark carts, and maintained darkness with high-volume white noise in the experiment room to minimize external stimuli. The water-restricted mice received water rewards 2.25 m down a 3-m visual virtual track, a track length comparable to other physical and virtual track place cell studies^3–7,30,31,34–39^. Mice were trained until they reached a behavioral task engagement level based on anticipatory slowing behavior before the reward (average 20±3.6 training days, 77.5±2.5 traversal laps per session; Fig. 1d-e; Extended Data Fig. 1; see Methods) comparable with other studies^35,36,38,39^. Once this criterion was reached, we used two-photon (2P) microscopy to image calcium transients in populations of dorsal CA1 neurons using GCaMP8m (Fig. 1c)^28–31,40^.

We imaged the same populations of neurons for five consecutive days. Since previous reports have suggested that technical difficulties in identifying and recording from the same neurons over days could explain previous observations of representational drift^41,42^, we developed a quantitative method based on volumetric plane registration to precisely identify the same imaging fields across days (Fig. 1f; see Methods). Our plane registration precision was ~2 µm, i.e., less than 15% of the diameter of an average mouse CA1 pyramidal cell (~13.2 µm ^43^). This robust registration methodology resulted in tight cross-correlations of the spatial footprints (Fig. 1g) and high correlations of cellular morphologies across days (median Pearson’s correlation of spatial footprints, r = 0.92, n = 8872 region of interest (ROI) pairs from 1351 cells active on multiple days, Fig.1h). We further visually confirmed that individual neurons were reliably identified based on their subcellular morphologies and were able to clearly identify the same processes across days (Fig. 1i-j, Extended Data Fig. 2a-b; see Methods).

A total of 1678 CA1 pyramidal cells in six mice were recorded (280±9.2 cells per mouse; Fig. 1i-j). On the first day, we identified 391 place cells based on spatial information^44^ (65.2±18.8 place cells/imaging field) and similar to previous reports^2–6,9,25,26^, observed a range of spatial tuning stability over the days of imaging, with some place cells expressing stable and others unstable tuning (Fig. 1j, Extended Data Fig. 2a-b). While the overall quality of the spatial code within days was preserved (Extended Data Fig. 2c, n = 6 mice, 2-way ANOVA across animals and days; total active neurons F(4,20)=0.18, p=0.95; fraction of place cells, F=0.33, p=0.85; spatial information of place cells, F=0.72, p=0.59), we observed qualitative differences in the population spatial tuning over the five days when we inspected sorted and cross-validated heat maps of the same cells (Fig. 1k). We then quantified representational drift with multiple statistical measures and found statistically significant drift over the five recording days in all eight measures (Fig. 1l-n; Extended Data Fig. 2d-e; 2-way ANOVA across animal and day, day effect; recurrence probability, F(4,20)=24.5, p~0; population vector (PV) correlation, F=22.3, p=3.9e-7; tuning curve (TC) correlation, F=4.8, p=0.0072; place field peak shift, F =5.9, p=0.0027; Bayesian decoding error, F=6.0, p=0.0024; Frobenius norm, F=37.2, p=5.3e-9; representational similarity, F=13.9, p=1.4e-5; representational drift index, F=21.9, p~0; see Methods; see Extended Data Table 1 for all ANOVA statistics). Our recurrence probability analysis revealed that only 164 out of the 391 place cells (41.9%) identified on Day 1 had the same place fields on Day 5. Therefore, significant representational drift occurs in the hippocampus of mice running on a linear treadmill in a familiar virtual environment, consistent with previous studies in virtual and real-world experimental settings^3–6^.

### Hippocampal representational drift persists even with highly reproducible behavior

Hippocampal place coding in mice shows dependencies on different behavioral factors, such as running speed and head direction^13–16^. Therefore, we asked whether differences in mouse behavior across days could explain drift in our experimental setup. We focused on the speed vs track position profiles for each traversal lap since this was one of the most salient behavioral differences across laps in our one-dimensional navigation task. For each mouse, from the dozens of laps per day, we sub-selected two different sets of laps. First, we sub-selected and found 20 laps that were both highly similar within each day and highly similar between days (“similar” set; Fig. 2a,c; Extended Data Fig. 3a; see Methods). Second, we sub-selected and found 20 laps that were highly similar within each day but were highly dissimilar between days (“dissimilar” set; Fig. 2b,d; Extended Data Fig. 3a-c). The within-day behavioral correlation was indistinguishable between the two sets (Fig. 2e; 3-way ANOVA across all laps, similar and dissimilar sets, animals, and days; set: F(2,78)=21.5, p=3.7e-8; Tukey-Kramer post-hoc test p = 0.61), while the across-day behavioral difference was significantly larger in the dissimilar sets (Fig. 2f; ANOVA across all days, similar and dissimilar sets and animals; similar and dissimilar sets: F(2,10)=190.1, p=1.1e-8; Tukey-Kramer post-hoc tests p<0.05).

**Figure 2:**
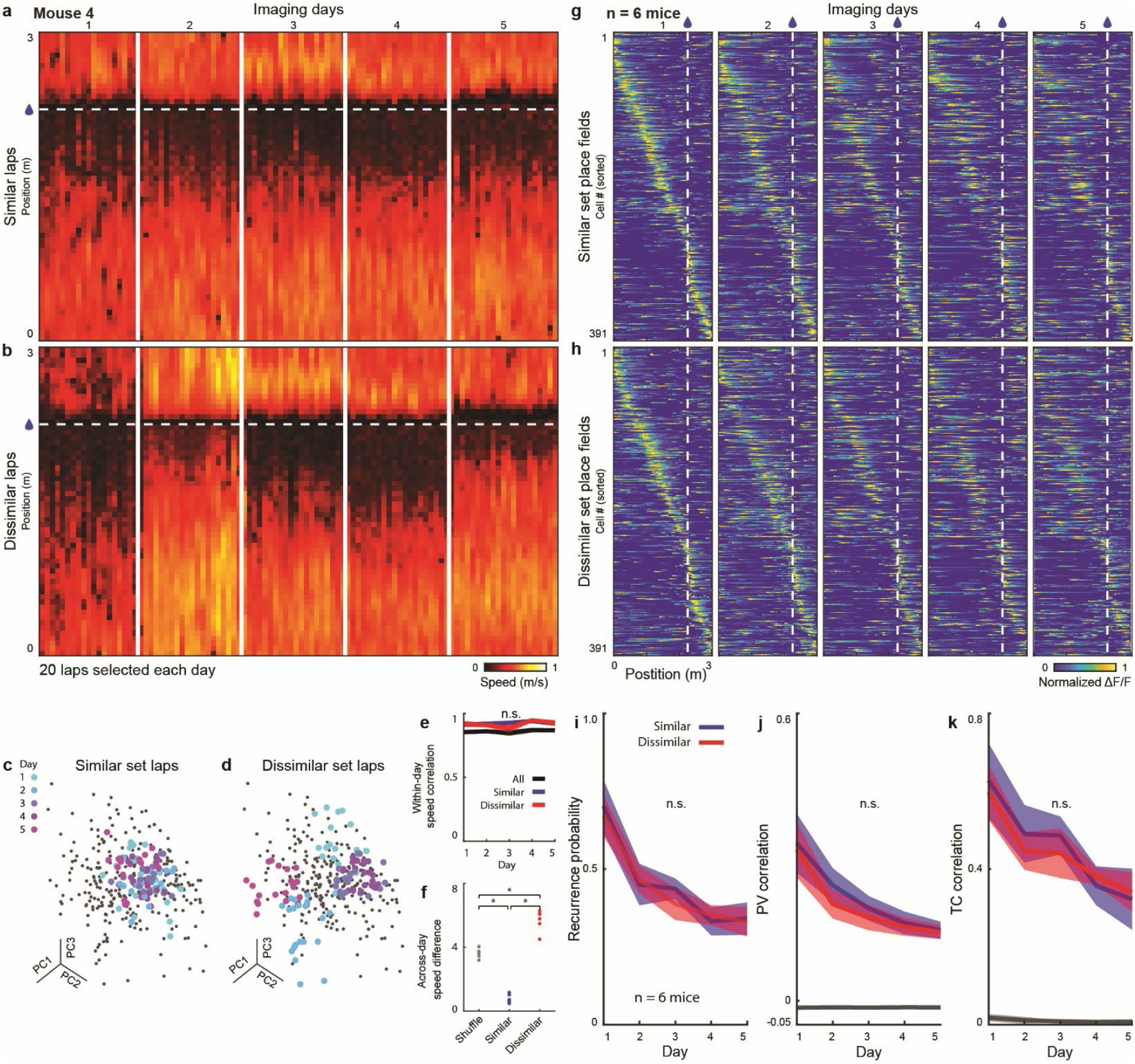
Hippocampal representational drift persists even with highly reproducible behavior. a-b) The speed profile of traversal laps of an example mouse across days, selected based on “similar” (a) or “dissimilar” (b) lap sets. 20 laps for each imaging day are selected for each subset. c-d) Principal component analysis (PCA) space representation of the laps (position binned speed vectors) of the example mouse in a-b for the “similar” (c) and “dissimilar” (d) lap sets. Each dot represents a single lap speed vector, and colors show different days. e-f) Within-day speed vector correlations for each of the five days (e) and across-day PCA space distance (f) (N= 6 mice, Tukey-Kramer post-hoc test, *p < 0.05). g-h) Cross-validated heatmaps of 391 cells (6 mice) identified as place cells on Day 1, plotted on similar and dissimilar laps. Day 1: the rows are sorted based on the place field locations across all laps, and only the similar (g) or dissimilar (h) sets mean DF/F is plotted. Days 2-5: same 391 cells sorted according to Day 1 order, plotting mean DF/F for similar (g) or dissimilar (h) lap sets on each indicated day. i-k) Representational drift measured by recurrence probability (i), population vector (PV) correlation (j), and tuning curve (TC) correlation (k) (i-k, 3-way ANOVA, sets effect, n.s. p>0.05). The shaded region shows SEM.

Since running speed was recently related to representational drift in mouse V1^20^, and flight path differences to drift in bat CA1^21^, we expected less drift when examining similar sets compared to dissimilar sets. However, when we examined the cross-validated place cell population heat maps restricted to the two different traversal sets, we observed no qualitative differences (Fig. 2g,h). Likewise, when we quantified representational drift using the multiple statistical measures described above, we found no statistical difference between the two sets (Fig. 2i-k, Extended Data Fig. 3d; 3-way ANOVA across similar and dissimilar sets, animals, and days, sets effect: recurrence probability F(1,49) =0.077, p=0.78; PV correlation F=1.6, p=0.22; TC correlation F=0.097, p=0.76; place field peak shift F=0.32, p=0.57; Bayesian decoding error F=2.0 p=0.16; Frobenius norm F=0.83, p=0.37; representational similarity F=0.72, p=0.40; representational drift index F=0.088, p=0.77). Therefore, the CA1 representational drift rate in mice is not detectably changed by differences in running behavior, with significant drift still occurring even when laps with highly similar behavior were selected across days.

Cognitive factors such as attention—which is often inferred from task engagement—could influence the rate of representational drift^36^. Previous studies have used pre-reward licking as evidence that mice have learned the location of a reward, and once learned, pre-lick serves as a measure of task engagement^4,35–39,47^. Comparable with these studies, our mice consistently and robustly exhibited pre-reward licking, indicating strong task engagement (Extended Data Fig. 1d-e and 5a-f). To investigate whether subtle differences in task engagement affect the rate of representational drift, we divided laps from each session into higher and lower pre-reward lick selectivity sets and found little to no significant difference in drift rates (Extended Data Fig. 4; 3-way ANOVA across days, animals, and set; set effect: recurrence probability, F(1,40) = 5.7, p=0.022; PV Correlation, F=0.87, p=0.36; TC correlation, F=5.5, p=0.024; place field peak shift F=1.2, p=0.27; Bayesian decoding error, F=7.2, p=0.011; Frobenius norm, F=2.3, p=0.14; representational similarity, F=5.0, p=0.031; representational drift index, F=2.1 p=0.15; see Methods). Furthermore, across all mice, we detected no significant correlations between average lick selectivity and drift rate in six of the drift metrics, weak correlation in one (Frobenius norm) and some correlation in one (representational similarity, Extended Data Fig. 8). Therefore, the CA1 representational drift rate in mice shows little or no change due to subtle differences in task engagement, with significant drift still occurring even in laps and mice with the highest lick selectivity.

### Hippocampal representational drift persists even in a highly reproducible sensory environment

Next, we hypothesized that subtle changes in the sensory environment over days might be the source of CA1 representational drift. Our visual virtual environment was the same each day and any auditory cues were masked with a white noise sound generator. In contrast, olfactory cues were not controlled in our (Fig. 1) or previous mouse experiments where significant hippocampal drift was observed^2–6,9,25,26^. Mice are highly olfactory animals, with the ability to detect extremely small variations in olfactory stimuli^17–19^. For example, mice display sensitivity and behavioral responses to odor concentrations^48^ below 1 × 10^−10^ M. Moreover, changes in the odor of an environment can induce changes in spatial tuning in hippocampal place cells^22,23,49–52^. Therefore, even small changes in olfactory cues from day to day could be perceptible to mice and could lead to differences in spatial representations over days.

To test the hypothesis that subtle odor variabilities may affect representational drift, we precisely controlled both the visual and the olfactory sensory cues available to the mice using our recently developed visual-olfactory multisensory VR system (Fig. 3a)^22,23^. We began with the “flat odor” task in which mice (n=5) were provided with a fixed concentration of α-pinene odorant via an airstream through a nose cone that enveloped their nose (Fig. 3b). During all training and imaging days, mice consistently experienced this odor identity and concentration. The visual VR environment was the same as used in Fig. 1 and, importantly, the within-day similarity between speed vs track position profiles for the flat odor group was indistinguishable from those of the “uncontrolled odor” group (Fig. 3e-f; Extended Data Fig. 1b-e, 5a-b; repeated-measures ANOVA on within-day correlation, task by day interaction; F(4,36)=0.49, p=0.74; across-day behavioral difference rank sum p=0.66). When we looked at the spatial code of CA1 place cells in the uncontrolled odor and flat odor tasks, we found no difference in the fraction of place cells recorded (Fig. 3g, 34.0% versus 36.2% in the uncontrolled odor versus flat odor task, χ^2^=0.70, p=0.40) but notably, there was a significant increase in the spatial information in place cells in the flat odor task (Fig 3h, rank-sum p=4.5e-6).

**Figure 3:**
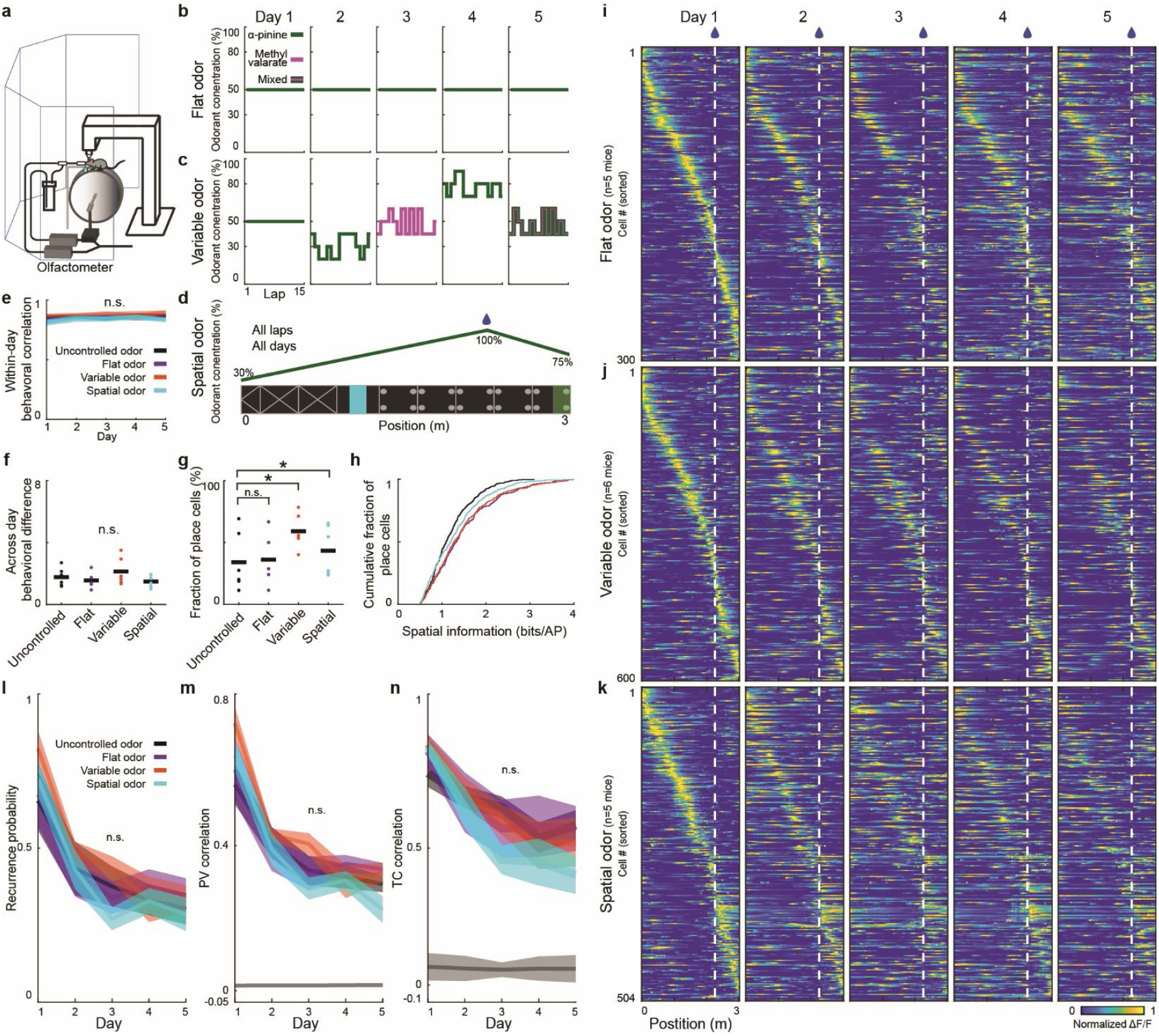
Hippocampal representational drift persists even in a highly reproducible sensory environment. a) Visual-olfactory multisensory VR provides a controlled odorant through a nose chamber while the mouse navigates the virtual environment. b-d) Schematics of odor presentation protocols across days. b) In the flat odor task, 50% α-pinene is presented on each lap. c) In the variable odor task, different concentrations and mixtures of α-pinene and Methyl valerate odorants were presented on each day. d) In the spatial odor task, α-pinene was presented with a fixed gradient profile across all laps and days. e) Within-day speed vector correlations for each of the five days in different tasks. f) Across-day PCA space distance in the different tasks (e,f, repeated-measures ANOVA, n.s., p>0.05). g-h) Fraction of place cells and their spatial information across different tasks. i-k) Cross-validated heatmaps of flat odor (i), variable odor (j), and spatial odor (k) tasks. l-n) Population representational drift in different odor tasks measured by recurrence probability (l) PV correlation (m), and TC correlation (n) (l-n, repeated measures ANOVA, task by day interaction, n.s. p>0.05).

Based on the previous odor-sensitive place cell studies^22,23,49–51^, we expected less drift in the flat odor group in comparison to the uncontrolled odor group. However, when we looked at the cross-validated place cell population heat maps, we observed no qualitative differences between these two groups (Fig. 1k, 3i). Similarly, when we quantified representational drift using the multiple statistical measures described above, we found no significant differences (Fig. 3l-n, Extended Data Fig. 5g; repeated measures ANOVA, task by day interaction; recurrence probability, F(4,36) =0.49, p=0.75; PV correlation, F=0.45, p=0.77; TC correlation, F=0.91, p=0.47; peak shift, F=1.56, p=0.21; Bayesian decoding error, F=0.42, p=0.79; Frobenius norm, F =0.28, p=0.89; representational similarity, F=0.88, p=0.49; representational drift index, F=1.13, p=0.36). Therefore, the CA1 representational drift rate in mice is not significantly changed when the sensory environment is precisely reproduced and controlled.

While drastic changes to the visual environment are known to cause partial and global remapping^53–55^, the impact of more subtle environment variability on representational drift rate across days is poorly understood. Therefore, to further test the effect of odor environment variability on representational drift rate, we next developed a “variable odor” task in which we used our multisensory VR system to systematically introduce different odors (α-pinene, methyl valerate, or mix) and concentrations across laps and days (Fig. 3c, Extended Data Fig. 6; see Methods). Again, the visual VR environment was the same as used in Fig. 1, and the within-day similarity between speed profiles along the track for this variable odor group (n=6) was indistinguishable from those of the uncontrolled and flat odor groups (Fig. 3c,e, Extended Data Fig. 5c-d, repeated-measures ANOVA on within-day similarity, task by day interaction, F(8,56)=0.43, p=0.90; ANOVA across-day behavioral difference, F(2,14)=0.95, p=0.41). Interestingly, the presence of the variable odor enhanced the CA1 spatial code, with a higher fraction of place cells in this task (Fig. 3g; 59.0%, a 1.7-fold increase compared to the uncontrolled task, χ^2^=144, p~0). Moreover, the place cells had higher spatial information than those in the uncontrolled odor task (rank-sum p=1.6e-6) and were similar to those in the flat odor task (p=0.58). When we examined the heat maps (Fig. 3j) and quantified representational drift rate using the multiple statistical measures, we found little to no difference between the three tasks (Fig. 3l-n; Extended Data Fig. 5g; repeated measures ANOVA, task by day interaction; recurrence probability, F(8,56) =2.0, p=0.068; PV correlation, F=2.1, p=0.046; TC correlation, F=1.1, p=0.41; place field peak shift F=1.4, p=0.23; Bayesian decoding error, F=2.6, p=0.018; Frobenius norm, F=1.7, p=0.11, representational similarity, F=1.3, p=0.25; representational drift index, F=0.87, p=0.55).

In addition, based on studies that show multisensory spatial information increases attention^2,4,56–67^, we hypothesized that enrichment of multisensory spatial information could change drift rate. Therefore, we designed a multisensory task in which both the visual and olfactory cues were spatially tuned and provided information about reward location (“spatial odor” task; Fig. 3d; Extended Data Fig. 5e-f; see Methods). The odor gradient peaked at the reward location and tapered off afterward, providing mice enriched multisensory information to integrate while navigating to the reward. The hippocampal drift rate of mice in this task was not different from the other three tasks (Extended Data Fig. 5g; repeated measures ANOVA, task by day interaction: recurrence probability, F(12,76) =1.6, p=0.11; PV correlation, F=1.8, p=0.062; TC correlation, F=1.6, p=0.11; place field peak shift, F=1.4, p=0.19; Bayesian decoding error, F=2.2, p=0.019; Frobenius norm, F=1.6, p=0.10; representational similarity F=1.9, p=0.049; representational drift index, F=1.62, p=0.10).

Finally, we asked whether similar variability as the variable odor task in the visual sensory modality could change the representational drift rate across days. We therefore designed a “variable visual” task where the virtual environment’s brightness varied lap-by-lap at different levels within and across days (Extended Data Fig. 7a-b; see Methods). We altered the overall brightness of the entire environment since such brightness changes also occur in natural habitats of mice, where variations in daylight (e.g., morning, noon, evening) and weather conditions (e.g., sunny or cloudy weather) create fluctuations in visual sensory input within and across days. The mice detected these brightness differences, indicated by their pupil size changes, with behavior comparable to the non-variable visual task (Extended Data Fig. 7c-f, within-day behavioral correlation, task by day interaction F(4,44)=0.19, p=0.94; across-days behavioral difference rank-sum, p=0.95). Notably, hippocampal drift rates under this visual variability task showed no statistical difference from the non-variable visual task (Extended Data Figure 7g-h, repeated measures ANOVA task by day interaction: recurrence probability F(4,44)=0.50, p=0.73; PV correlation F=0.38, p=0.82, TC correlation F=0.49, p=0.74; peak-shift F=0.74 p=0.57; Bayesian decoding error F=0.049, p=0.99; Frobenius norm F=0.34, p=0.85; representational similarity F=0.77 p=0.55, representational drift index F=1.12, p=0.36). Therefore, overall, the CA1 representational drift rate in mice is not detectably changed by differences in sensory environment variability across days, with similar significant drift still occurring when the sensory environment is either precisely controlled or more varied.

Our analysis so far has focused on place cells encoding track position, but it is possible that other variables are also significantly encoded, or influencing the spatial coding, during our tasks^68^. To examine these possibilities, we first constructed “internal tuning curves”^69^ based solely on population activity, independent of the mouse’s track position. For neurons with internal tuning curves, we (separately) applied our original spatial position analysis and compared the internal and spatial tuning curves across the population within each session. We found that the two tuning curves were highly similar, with the overall population structure highly similar between the two forms of analysis (Extended Data Fig. 9). Second, based on a recent study showing that hippocampal representations can be influenced by the sensory input associated with each human experimenter’s identity^70^, we sought to determine whether the experimenter(s) could influence drift rates. We ensured that a single experimenter handled each mouse in all five imaging days across our experiments. We then examined whether different experimenter identities were associated with different rates of representational drift, but found no significant effect of experimenter identity (n=2 experimenters) on the drift rate (Repeated measures ANOVA, experimenter by day interaction: recurrence probability, F(4,108) = 0.83, p=0.51; TC correlation F=1.14, p=0.34; PV correlation F=1.07, p=0.38; place field peak shift F=1.6 p=0.17; Bayesian decoding error F=0.37 p=0.82;

Frobenius norm, F=1.63 p=0.17; representational similarity, F=1.00, p=0.41; representational drift index, F=1.55, p=0.19). Thus, consistent with previous work^69^, we conclude that space (track position) is the dominant feature of the hippocampal representation during our tasks and experimenter identity did not significantly change drift rate.

### Higher neuronal excitability is predictive of more stable representation in hippocampal CA1

We have established that neither behavioral nor sensory variabilities in our tasks explain hippocampal CA1 representational drift in mice across days, suggesting that drift is not externally driven and may instead be determined by intrinsic cellular or circuit mechanisms. Though some functional properties of place cells have previously been associated with stability over days (little drift on the single cell level), such as lap-by-lap variability and place field distance to reward^36,71^, a systematic analysis of such properties as well as properties related to the recording signal quality that could impact cell detection over days has not been performed. Therefore, we divided place cells from the 30 mice across our five tasks into stable place cells (recurred for 4 or more days based on recurrence probability; 9.1%, 731/8014 of all neurons; 18.0% of place cells per mouse; see Methods) and unstable place cells (recurred for two or fewer days based on recurrence probability; 18.3%, 1463/8014 of all neurons; 36.0% of place cells per mouse; Fig. 4a-e, Extended Data Fig. 10, 11a; see Methods). The remaining ~46.0% of place cells (23.4%, 1868/8014 of all neurons) were excluded from the following analysis since they could not be clearly categorized as either stable or unstable. As expected from these definitions, heat maps of stable cells were highly similar over days and heat maps of unstable cells were highly dissimilar. These observations were quantified using various measures of representational drift (Fig. 4f-h, Extended Data Fig. 11b, n= 30 mice, sign rank tests between stable and unstable cells on emerge day+2, recurrence probability, p = 2.6e-6; PV correlation, p = 2.6e-6; TC correlation, p=2.6e-6; place field peak shift, p = 2.6e-6; Frobenius norm, p = 6e-5; representational similarity, p=2.6e-6).

**Figure 4:**
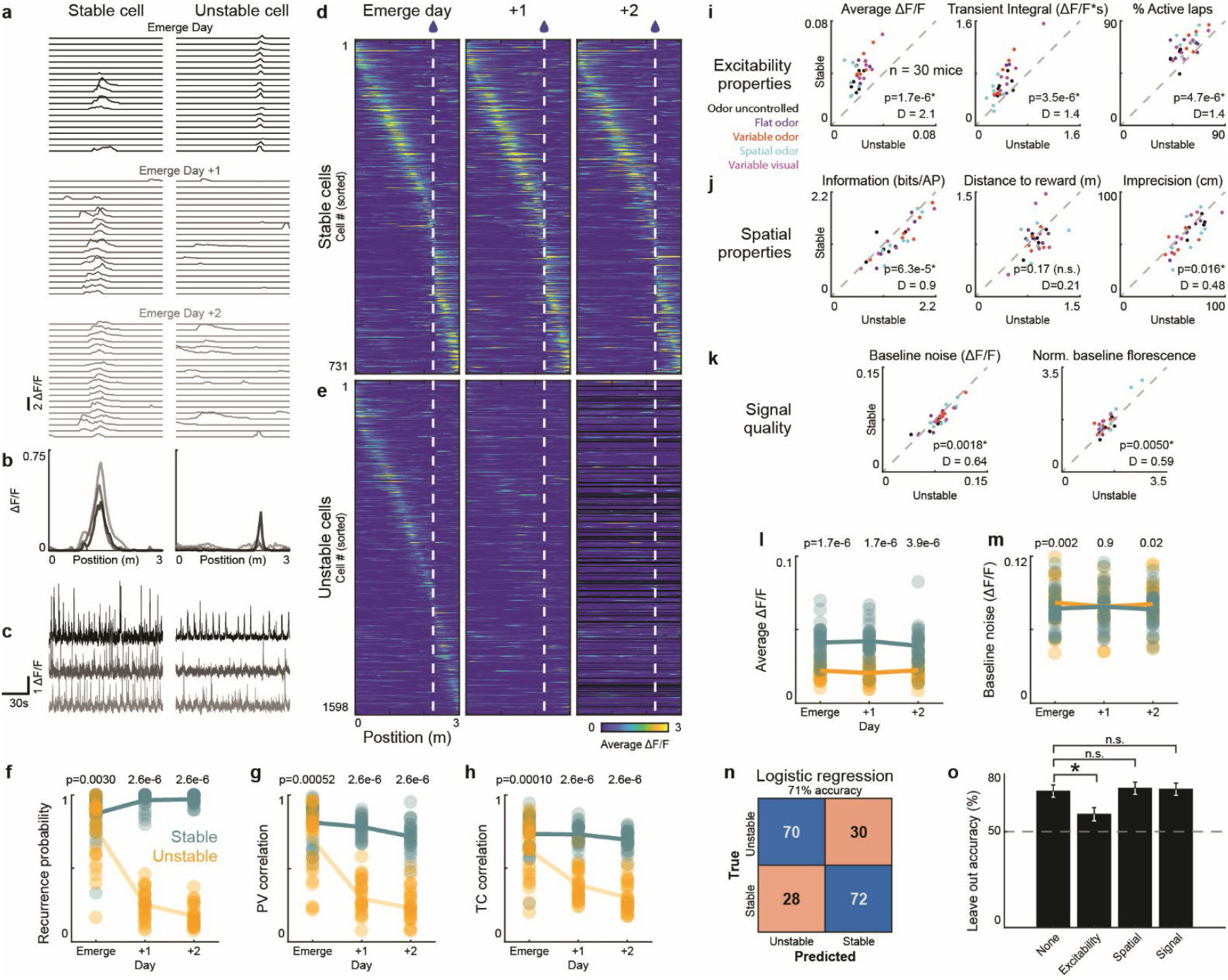
Higher neuronal excitability correlates with and is predictive of more stable representation in hippocampal CA1 place cells. a-c) Example stable and unstable place cells. Lap by lap DF/F vs position (a) and mean tuning curves (b) from “emerge day” to two days after and their corresponding DF/F vs time traces (c). d-e) Cross-validated heatmaps for all stable and unstable place cells from the emerge day and two following days. f-h) Representational drift of stable and unstable cell populations across days (each point represents a mouse, n=30 mice) measured by recurrence probability (f), PV correlation (g), and TC correlation (h) (f-h, sign rank test). i-k) Stable vs. unstable place cell excitability properties (i), spatial properties (j), and signal quality (k). Each point is the emerge day average for each mouse (n=30 mice, Wilcoxon signed rank tests and Cohen’s d). I-m) Average dF/F (l) and baseline noise (m) from the emerge day and two following days for stable and unstable cell populations (n=30 mice) (l,m, sign rank test). n) Logistic regression classification of stable vs. unstable place cells. The confusion matrix of the fitted regression model predicted whether a place cell becomes a stable or unstable place cell from the emerge day properties. o) Accuracy of the logistic regression prediction when each group of properties is left out. Error bars show 95% binomial confidence intervals; likelihood ratio (χ^2^) test.

Since CA1 drift does not appear to be externally driven, we examined whether any functional properties of place cells on the “emerge” day was correlated to stability over subsequent days. We found that several cellular features related to neuronal excitability of the stable cell groups were significantly greater with larger effect sizes, compared to those of the unstable cell groups on the emerge day (Fig. 4i; Extended Data Fig. 11c-d; Wilcoxon signed rank tests and Cohen’s d; average DF/F, p=1.7e-6, d=2.1; transient integral, p=3.5e-6, d=1.4; %active laps, p=4.7e-6, d=1.4; %significant transients, p=1.7e-6 d=2.1; TC peak, p=1.7e-6 d=1.6) and subsequent days (Fig. 4l, Extended Data Fig. 11g). In contrast, we found some significant differences, but with only small effect sizes, between stable vs unstable cell groups in their spatial tuning properties on the emerge day (Fig. 4j, Extended Data Fig. 11e, h; spatial information, p = 6.3e-5,d=0.90; place field distance to reward, p = 0.17, d= 0.21; place cell firing imprecision, p = 0.016, d = 0.48). Finally, as a control for properties related to our imaging quality, we examined signal quality measures such as baseline noise and baseline fluorescence on the emerge day (Fig. 4k; Extended Data Fig. 11f) and across days (Fig. 4m; Extended Data Fig. 11i) and only observed weak statistical differences with small effect sizes between stable vs unstable place cell groups (baseline noise, p = 0.0018, d = 0.64; baseline fluorescence, p = 0.0050, d = 0.59). These signal quality features also showed weak, if any, correlations with excitability and spatial measures of place cells (Extended Data Fig. 12). Lastly, when we extended the same analysis to all active cells (not just place cells) we found highly similar results (Extended Data Fig. 13; see Methods). Therefore, greater neuronal excitability, rather than spatial tuning properties or experimental recording differences, were most correlated to place field stability over the following days.

Based on these results, we then asked how well the excitability features on the emerge day could predict the destiny of place cells (i.e., binary classification of stable vs. unstable place cells) over days using elastic-net logistic regression analysis (Fig. 4n-o; see Methods). Our regression model, trained with multiple excitability, spatial and signal quality properties, was able to predict and classify place cells with 71% accuracy on a 200-neuron counter-balanced test data set (Fig. 4n). When excitability, spatial, or signal quality features were left out of the model training steps, only leaving out excitability properties had a significant impact on the prediction accuracy of place cell stability over days (Fig. 4o; excitability properties, χ^2^= 6.4, p=0.011; spatial properties, χ^2^= 0.11, p=0.74; signal quality, χ^2^= 0.049, p=0.82). Therefore, not only does neuronal excitability correlate with place field stability over days, but also this intrinsic property has significant power to predict which place fields are more likely to persist or drift.

## Discussion

Here, we were motivated by the question of whether behavioral and sensory variabilities could provide a general explanation for representational drift in the hippocampus. To answer this, we utilized our recently designed multisensory VR system and developed an online volumetric plane registration method that allowed us to precisely and robustly identify the same cells across days with ~2 um error in XYZ planes. This registration method represents a marked improvement over the across-day imaging plane identification methods used in previous representational drift research.

We first compared drift rates using laps with similar and dissimilar running speed profiles across days and found no detectable difference in the rates (Fig. 2). Our result supports the idea that hippocampal drift in mice is not caused by overt behavioral variability across days, though we cannot rule out the possibility that some subtler behavioral variables that we did not consider here might have varied across days and led to the observed drift. Our result differs from recent research describing the modulation of V1 representations by running behavior over days^20^ which suggests that the stability of the primary sensory region may be more heavily influenced by behavioral changes than higher association areas, such as the hippocampus. However, a recent hippocampal place cell study in bats^21^ found little or no drift when considering constrained flight paths with highly stereotyped speed patterns, which brings species differences into the mix of factors possibly influencing representational drift^41^. More research is clearly required to determine how behavioral influences on representational drift vary across brain regions and animal species.

Mouse behavior can be strongly impacted by even small changes in their sensory environment. Since olfactory cues are known to be involved in driving place firing in CA1^2,22,23,49,51,72,73^, we reasoned that a significant uncontrolled sensory feature in previous mouse CA1 representational drift studies^3–5,26^ was the olfactory environment, particularly surface odors which are difficult to clean and could change over days and be detected by the mice. However, we found that the CA1 representational drift rate was comparable between controlled (flat odor) and uncontrolled odor tasks. Further, even when we introduced variable odor (Fig. 3) and variable visual tasks within and across days (Extended Data Fig. 7), the drift rate was similar and did not increase. Our result supports the idea that hippocampal drift in mice is not caused by overt olfactory or visual variability across days. Though at first glance our result may seem surprising, we recently showed that only sensory modalities with behavioral relevance have a strong influence on the cognitive map^23^. This prior research did not examine such influences on the stability of the cognitive map over days, but when combined with our results here, we can speculate that representational drift is also not influenced by variable sensory modalities that have little behavioral relevance.

While it is unclear what role CA1 representational drift plays in memory and cognition, our results support the idea that drift is an internally generated phenomenon, rather than externally driven. We were able to determine several cellular features that predicted place cell stability (Fig. 4), which may help illuminate the neural mechanisms of drift and lead to a deeper understanding of drift’s cognitive role in the future. Our finding that excitability features of cells are predictors of their long-term stability (i.e., little drift) is particularly interesting in the context of previous research that has found a link between excitability and the likelihood that a CA1 neuron is recruited to encode a new experience^31,74–76^. Indeed, in computational research, simulated Hebbian networks with fluctuating excitability over time could generate representational drift^77^. Further, excitability is a key parameter in memory linking, with more excitable cells being more likely to link memories across contexts^78^ and, interestingly, a reduction in the excitability of hippocampal neurons occurs during aging^79^. Thus, our finding may imply that the hippocampal representational drift rate increases during aging, leading to a prediction that could be tested in future research.

While our results support the idea that drift is an internally generated phenomenon largely related to cellular excitability, this does not mean that external factors do not (or cannot) impact the rate of representational change. For example, as more and more sensory features of an environment are changed, smaller changes to the representation (e.g. rate remapping) are observed, and eventually, large changes occur (e.g. global remapping)^53,54^. Therefore, external factors certainly play a role in determining how consistent a representation is over days. Our study instead explores the other end of the spectrum: when external factors are highly stable or varied in a systematic way, how stable is the hippocampal representation? Our finding that representational drift still persists when external factors are clamped/controlled (as much as currently possible) and even when external factors are subtly changed indicates that, in such scenarios, the drift rate is largely driven by intrinsic factors for which intrinsic cellular excitability is the strongest determining factor that we identified.

We controlled for behavioral and sensory environment variability at comparable levels as the recent bat CA1 research, but still found CA1 representational drift. Therefore, our results might alternatively suggest that the CA1 regions of bats and mice are fundamentally different in terms of representational stability. This idea is further supported by previous research, which has found significant differences between these two species regarding hippocampal place cell properties and oscillations^80,81^ and cellular-level electrophysiological properties^82,83^. Importantly, it is currently unknown whether bats have overall more excitable CA1 neurons compared to mice and whether such potential differences in excitability between mice and bats lead to the observed differences in representational drift.

Representational drift has been observed across numerous brain regions, tasks and species, with rates of drift that vary considerably^21,41,84–89^. For example, neural representations are relatively stable in motor areas^41,90–92^ but not in the primary olfactory cortex^89^ or the posterior parietal cortex^87^. It is possible that differences in neuronal excitability may underlie many of these previously observed differences in drift rate. However, these previous studies also used vastly different behaviors, which may also explain differences in drift rate. For example, the highly stereotyped motor patterns exhibited by lever pressing mice^93^, flying bats^21^ and singing songbirds^41^, may be fundamentally different types of behavioral stereotypy compared to the behavioral similarity considered here (laps with more similar speed or pre-lick profiles). Moreover, because our experiments were performed in highly controlled VR conditions, they do not fully reflect the complexity of natural environments. Thus, it remains uncertain how well our findings will translate to more naturalistic settings, where animals experience a broader range of sensory input variability and complex behaviors^94–96^. Also, while we took extensive measures to minimize animals’ experiences outside the experiments (e.g., single housing, no cage changes, no running wheel, enclosed transport), we did not systematically monitor or control variables such as activity levels, sleep, grooming, or foraging behaviors in the home cage or other factors such as hippocampal replay occurring outside of experimental sessions^97,98^. In the end, more research is needed to determine how these and all the factors considered here fit into a general explanation for the drift phenomenon across brain regions and species.

## Extended Data Figures

**Extended Data Figure 1:**
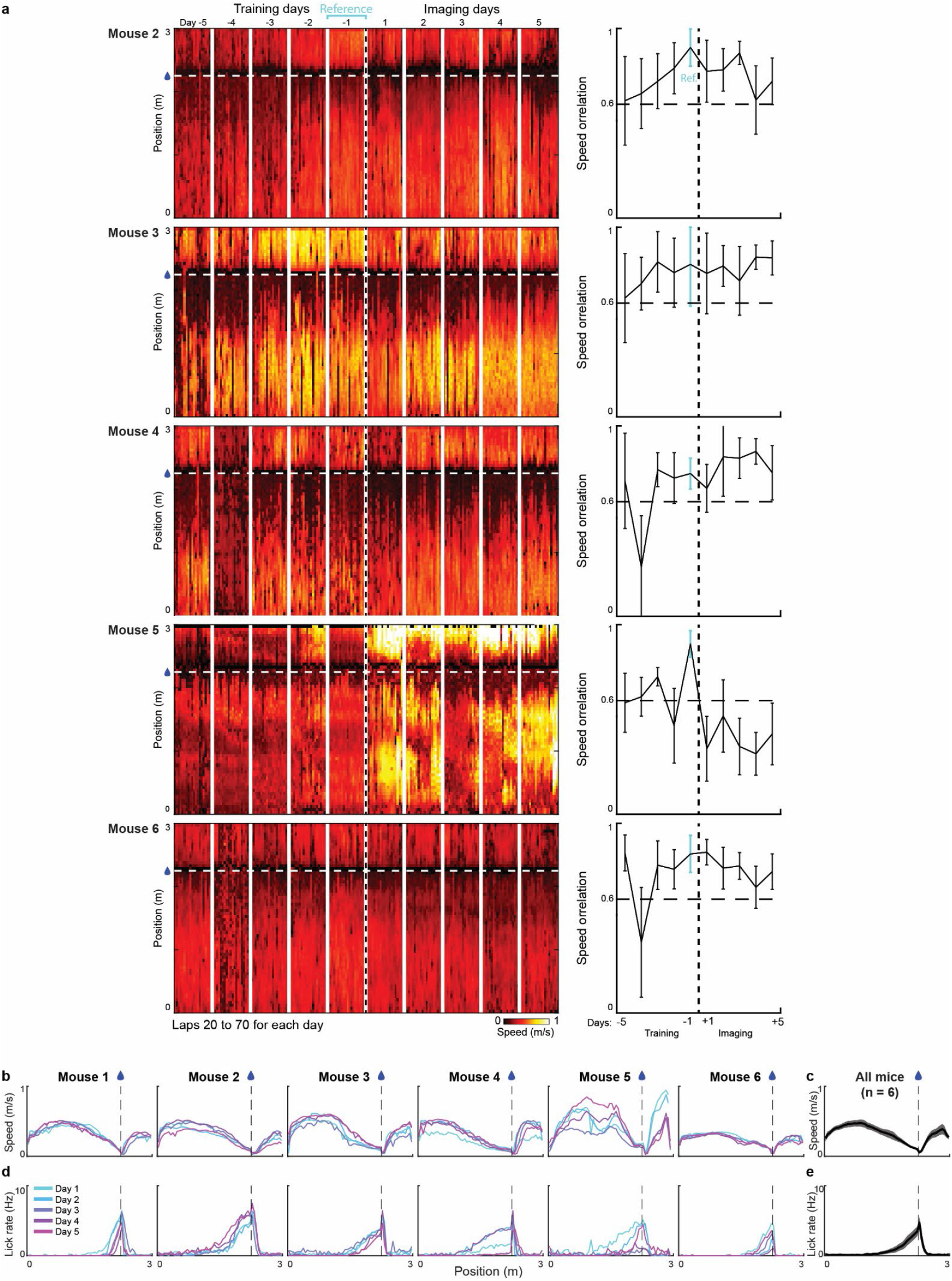
Running behavior of individual mice during training and imaging days. a) Left: The speed profile (m/s) of five mice performing the task in a familiar environment over five training and five imaging days. Each column shows a single-track traversal lap. The horizontal dash line indicates the water reward location, and the vertical white lines separate each day. Right: The speed correlation of laps 11-30 on each day to the mean lap from laps 11-30 on the reference day (cyan). Horizontal dashed line indicates the correlation threshold. b-c) The average speed profile of each mouse versus track position across days (b) and their mean (n=6 mice) (c). d-e) The average lick profile of each mouse versus track position across days (d) and their mean (n=6 mice) (e).

**Extended Data Figure 2:**
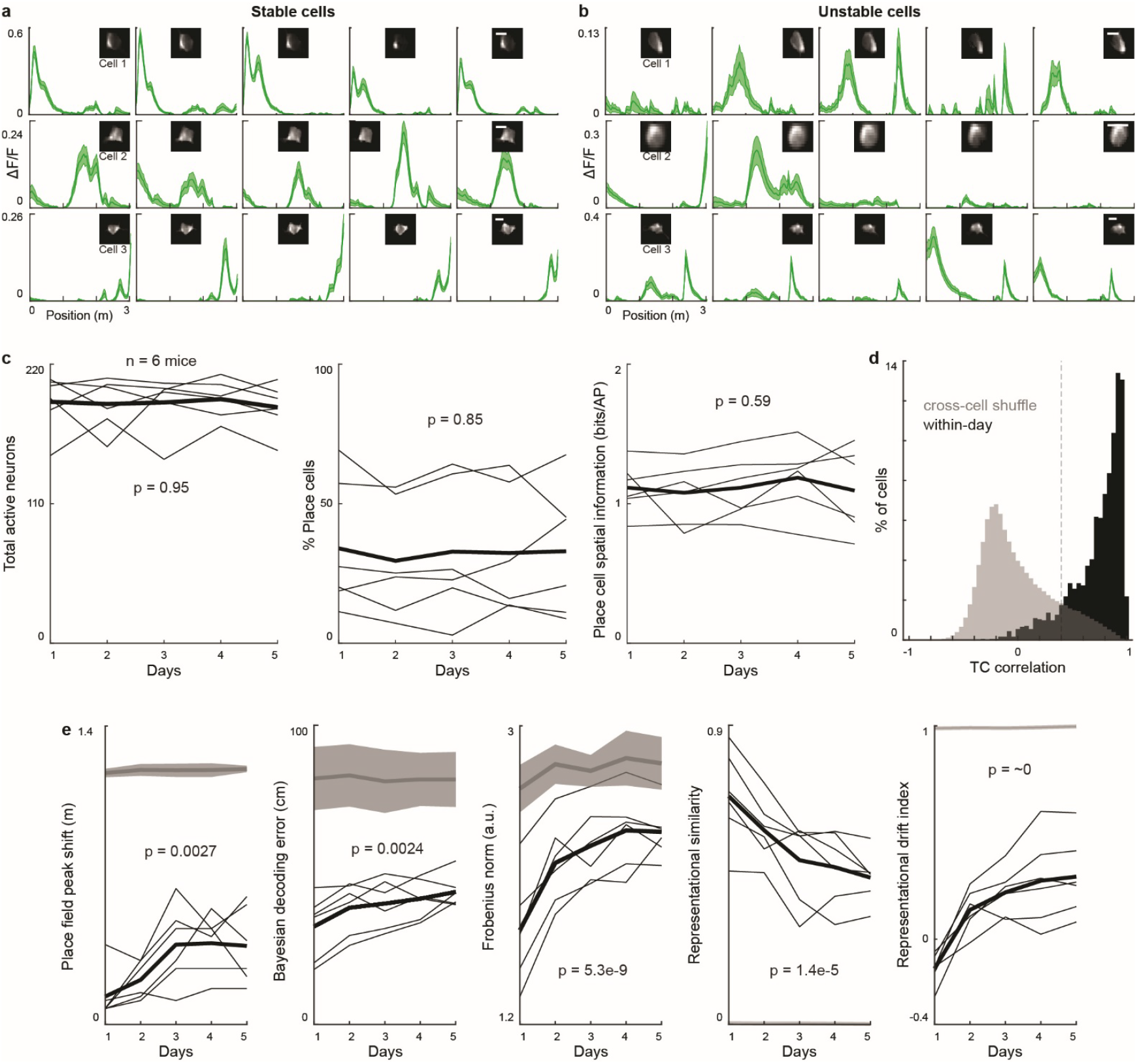
Cell type examples and additional representational drift measurements. a-b) Additional example cells showing stable (a) and unstable (b) spatial tunings across days. Scale bar=10 um. c) Quantification of total number of active neurons (left), fraction of active neurons that are place cells (middle), and spatial information of place cells (right), 2-way ANOVA across animals and days. The thick black line shows the average. d) Tuning curve (TC) correlations for place cells within day. Odd versus even laps (black) are separated from 10,000 randomly chosen pairs of place cells (gray) at 0.4 threshold (dashed line). e) Representational drift measured by place field peak shift, Bayesian decoding error, Frobenius norm, representational similarity, and representational drift index, 2-way ANOVA across animals and days.

**Extended Data Figure 3:**
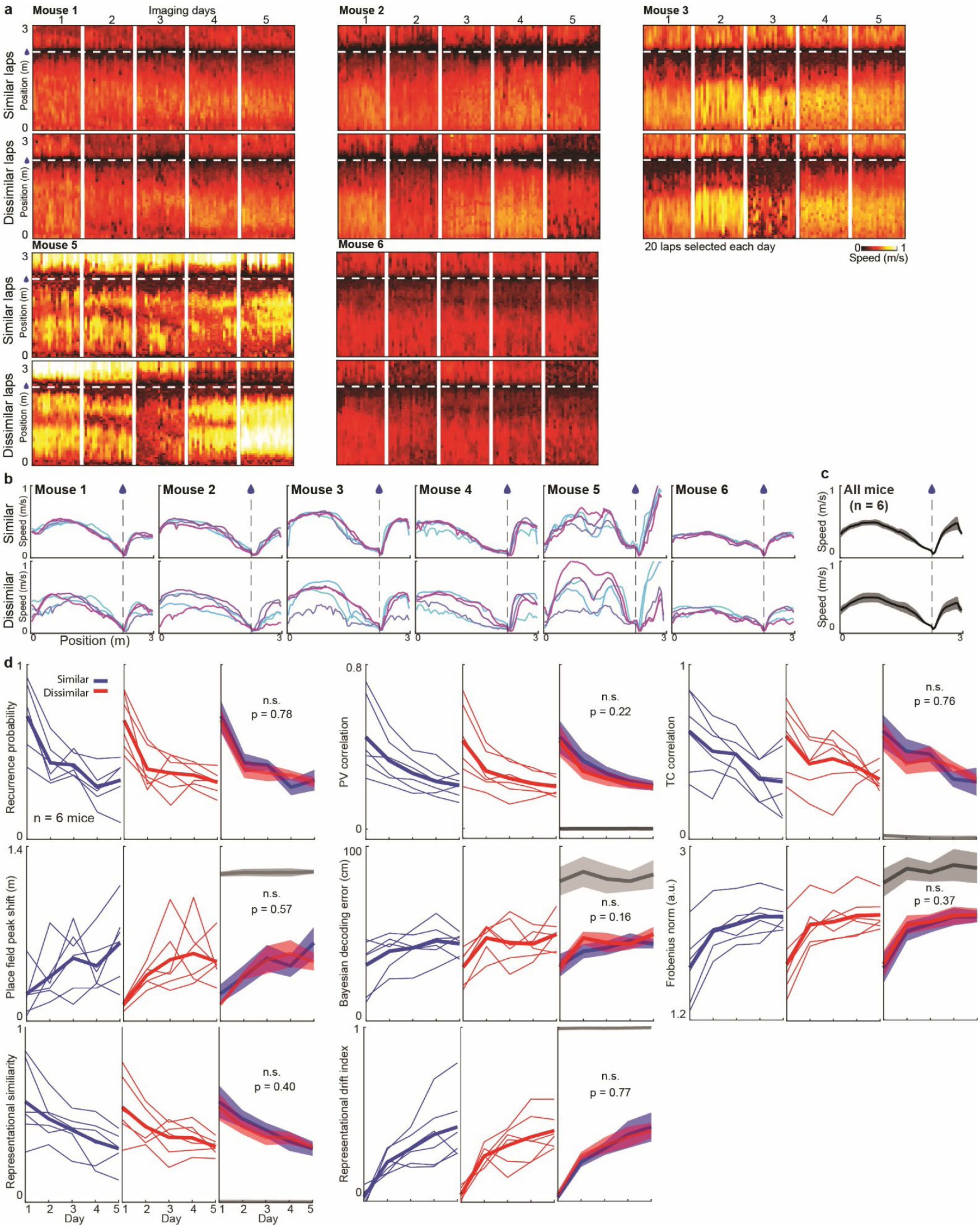
Running behavior and representational drift measurements for similar and dissimilar sets. a) The speed profile of mice across days; laps selected based on similar or dissimilar running sets. b-c) The average speed profile of the similar and dissimilar running laps of each mouse versus track position across days (b) and their mean (n=6 mice) (c). d) Representational drift of similar (blue) and dissimilar (red) running sets measured by various measurements (3-way ANOVA across animal, day and set; lap set effect, n.s. p>0.05) The shaded region shows SEM.

**Extended Data Figure 4:**
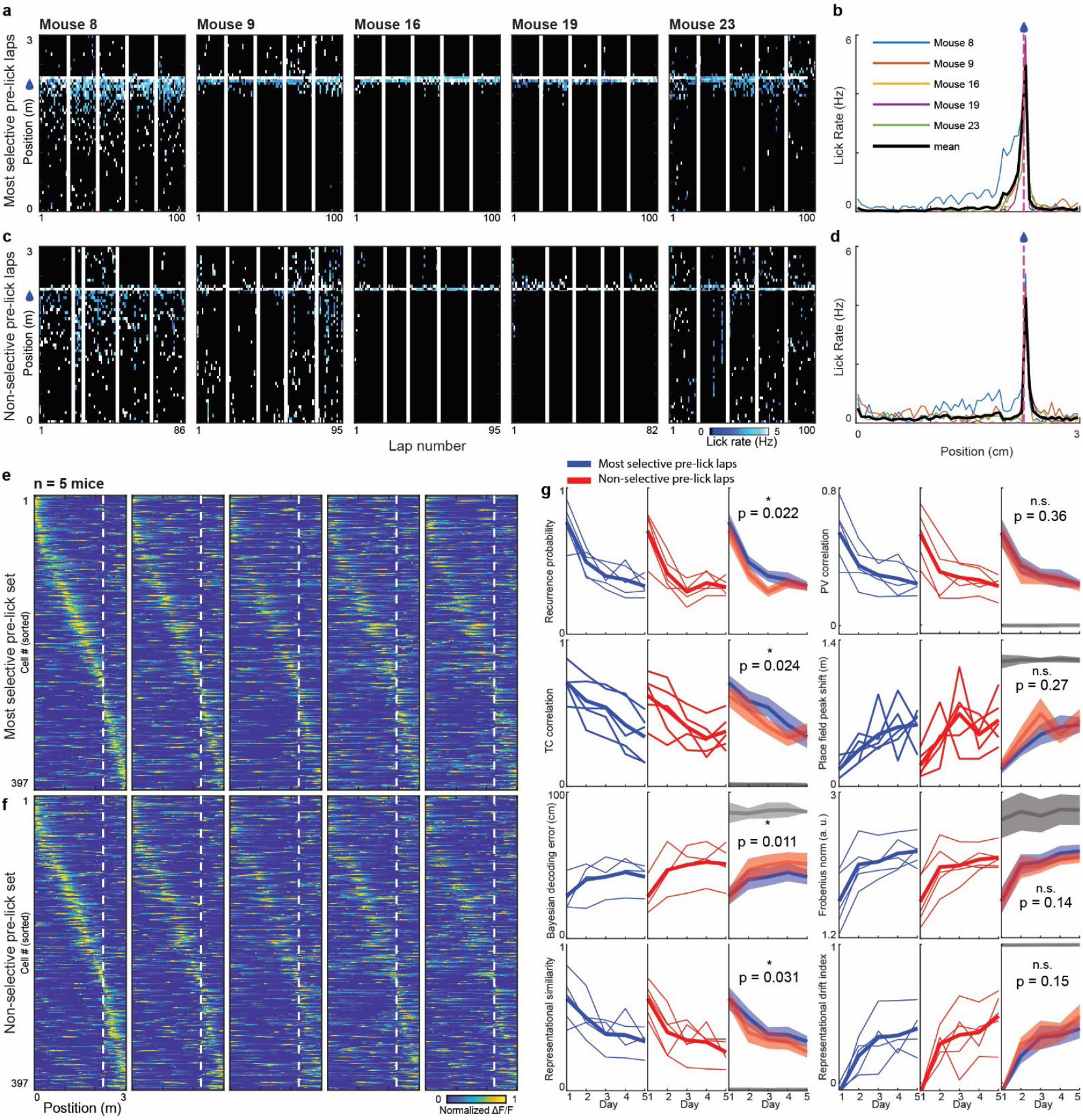
Licking behavior and representational drift measurements for most selective and non-selective pre-lick sets. a-d) The lick profile of mice across days for most selective (a-b) and non-selective (c-d) pre-lick laps. The tick black curve shows the average lick rate of all mice, and the red dashed line indicates reward location (b and d). e-f) Cross-validated heatmaps of 397 cells (6 mice) identified as place cells on Day 1, calculated based on most selective (e) and non-selective (f) pre-lick sets. g) Representational drift of most selective (blue) and non-selective (red) pre-lick sets measured by various measurements, 3-way ANOVA across days, animals, and set (lap set effect). The shaded region shows SEM.

**Extended Data Figure 5:**
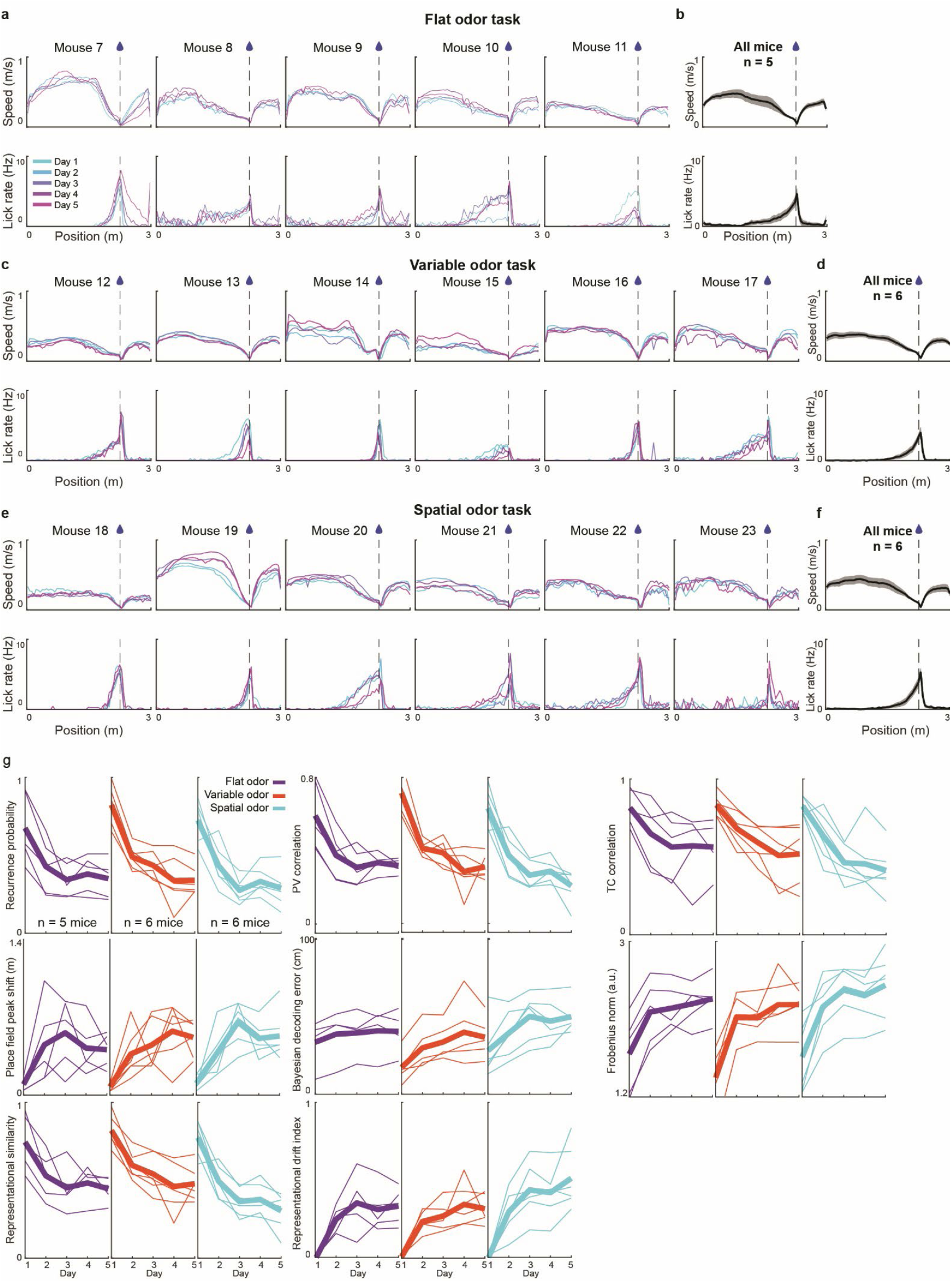
Running behavior and additional representational drift measurements for flat, variable, and spatial odor tasks. a-f) The average speed and lick profiles of each mouse versus track position across days in flat odor (a), variable odor (c), and spatial odor (e) tasks and their mean (b, d, and f). g) Representational drift of various odor mouse groups measured by various measurements. The shaded region shows SEM. The three groups have little or no statistical difference by multiple repeated measures ANOVA comparisons (see Extended Data Table 1 for all comparisons).

**Extended Data Figure 6:**
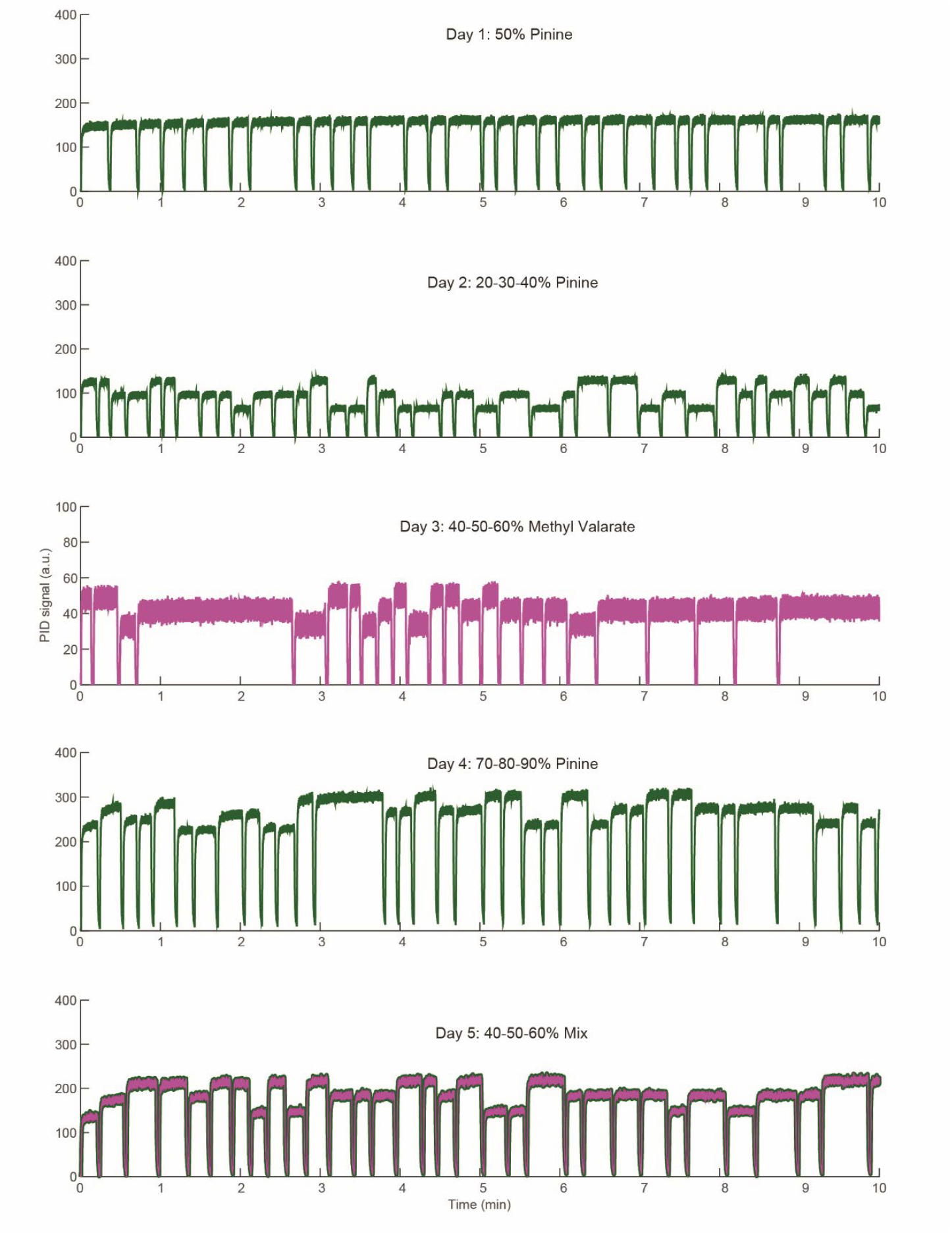
Odor concentration measurements for variable odor task using photoionization (PID) detector. PID measurements acquired from the olfactometer nose cone for the five-day protocol of the variable odor task described in Fig. 3c. Day 1 of the variable odor protocol is the same as the flat odor protocol in Fig. 3b. The measurements were performed while a mouse was running on a treadmill in a virtual environment. In the inter-trial-interval, after the mouse completed one lap and before the start of the next, the odor concentration was set to 0.

**Extended Data Figure 7:**
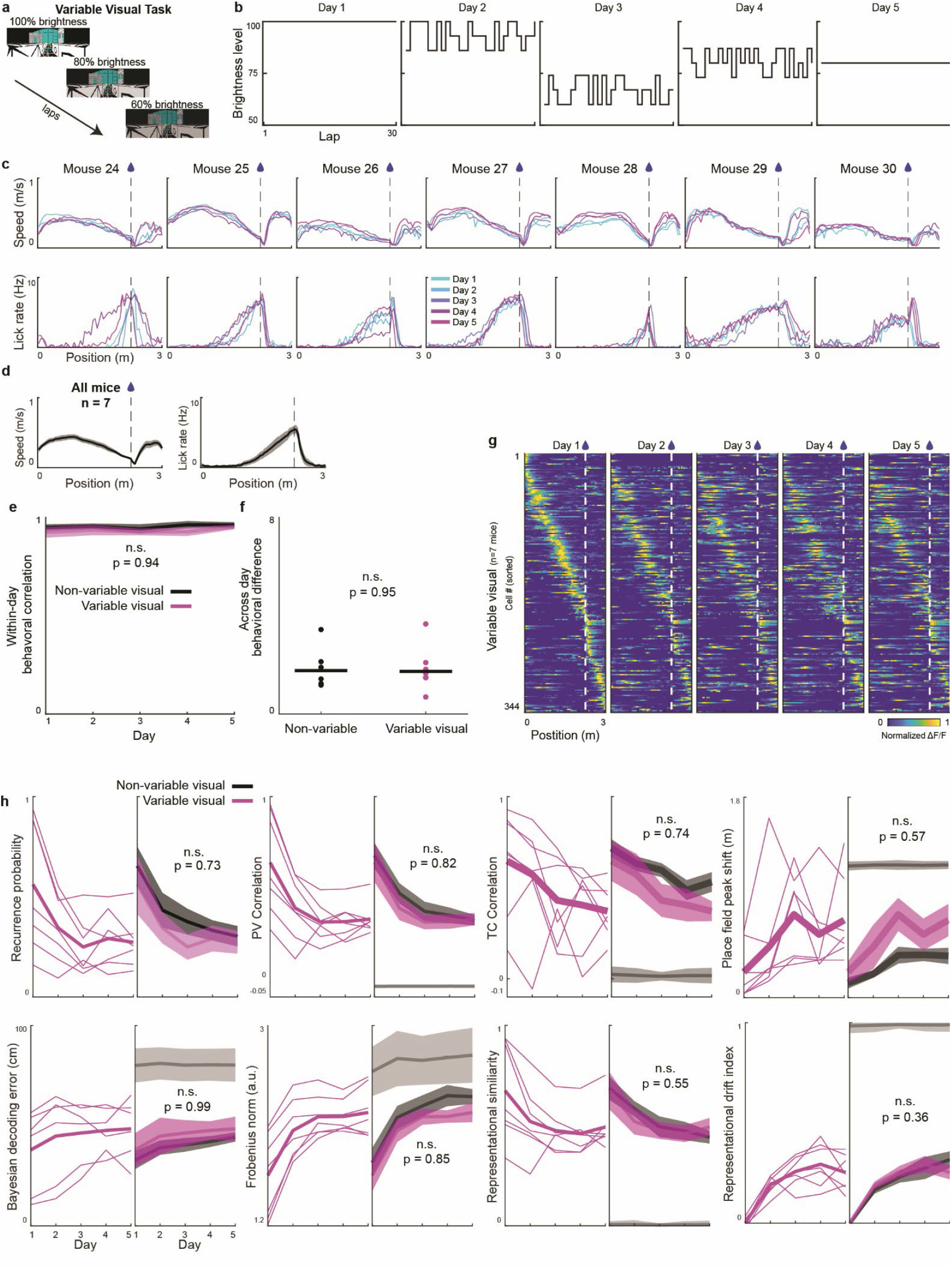
Subtle sensory visual variability does not detectably affect hippocampal representational drift in mice. a-b) In variable visual task, VR brightness varies lap-by-lap at different levels within and across days). c-d) Running and licking behavior of individual mice (c) used in the task and their averages (n=7) (d). e-f) Within-day speed vector correlations for each of the five days (e, task by day interaction) and across-day PCA space distance (f, rank-sum) (n=7 mice, n.s., p > 0.05). g) Cross-validated heatmaps of 344 cells (7 mice) identified as place cells on Day 1. h) Representational drift of variable visual (pink) and non-variable visual (black) mouse groups measured by various measurements, repeated measures ANOVA task by day interaction. The shaded region shows SEM.

**Extended Data Figure 8:**
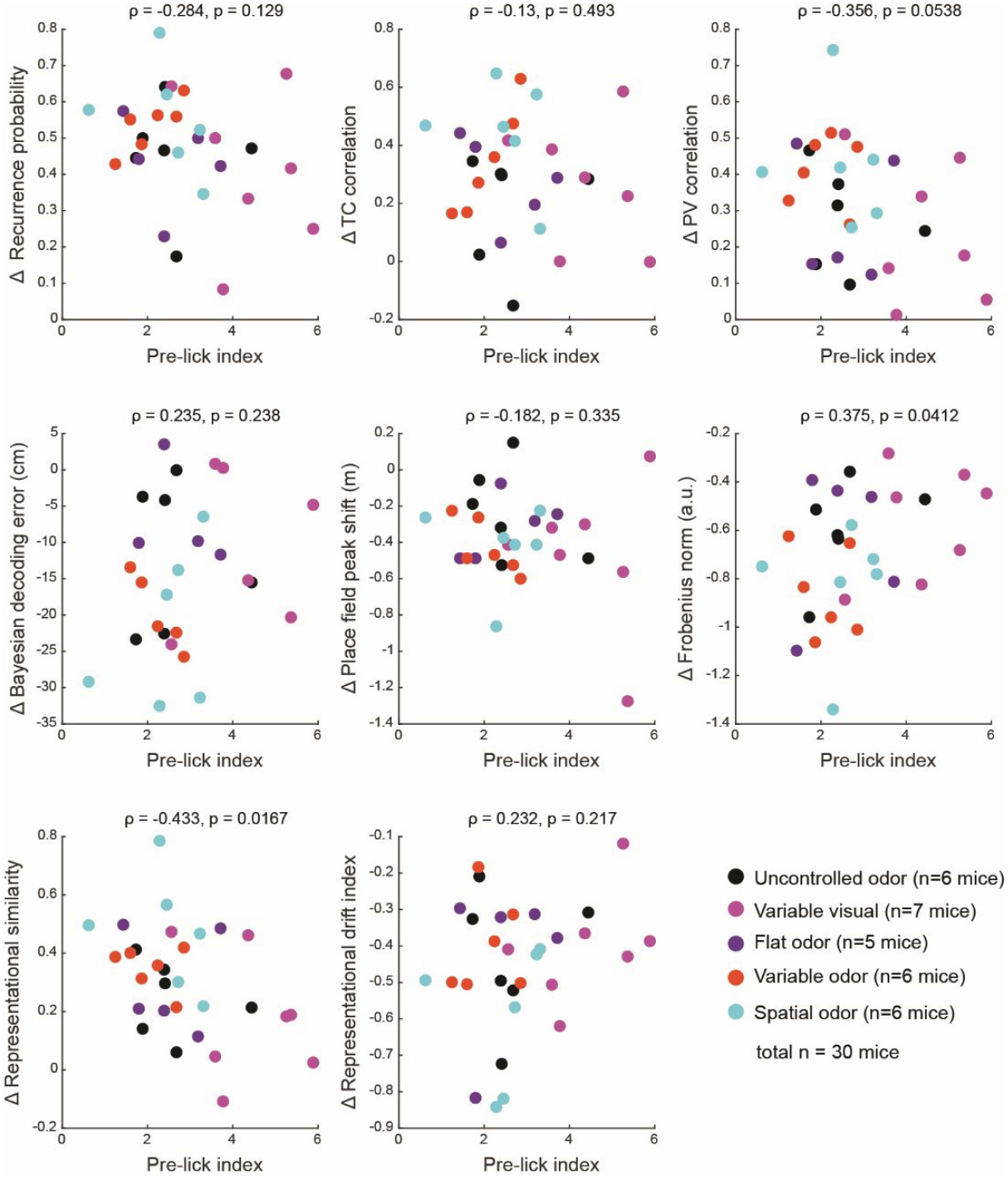
Plots of representational drift measurements vs pre-lick index across all mice. Plots of representational drift measurements vs pre-lick index across all mice (n=30 mice) for five different tasks. The x-axes are the average prelick index across all days and the y-axes are the difference between the first and last day values for each drift measurement. Pearson correlation ρ and p-values are listed above each panel.

**Extended Data Figure 9:**
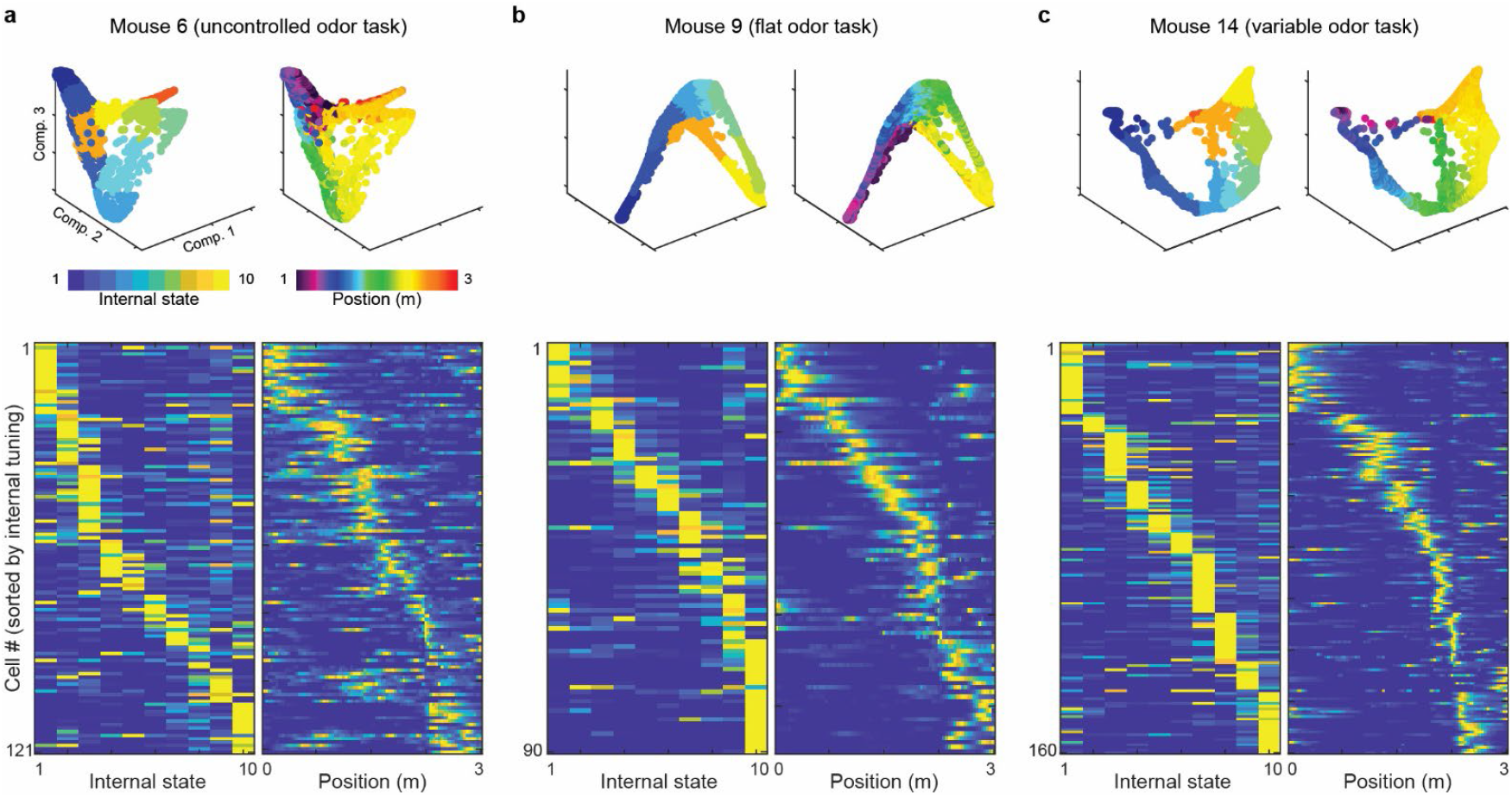
Examples of hippocampal internal states and spatial tunings. Internal and spatial tuning curves of hippocampal neuronal activities for example mice from uncontrolled odor (a), flat odor (b), and variable odor (c) tasks. Top: Internal and spatial tuning of hippocampal population activity in PCA space. Bottom: Cross-validated heatmaps of cells calculated based on internal state and position.

**Extended Data Figure 10:**
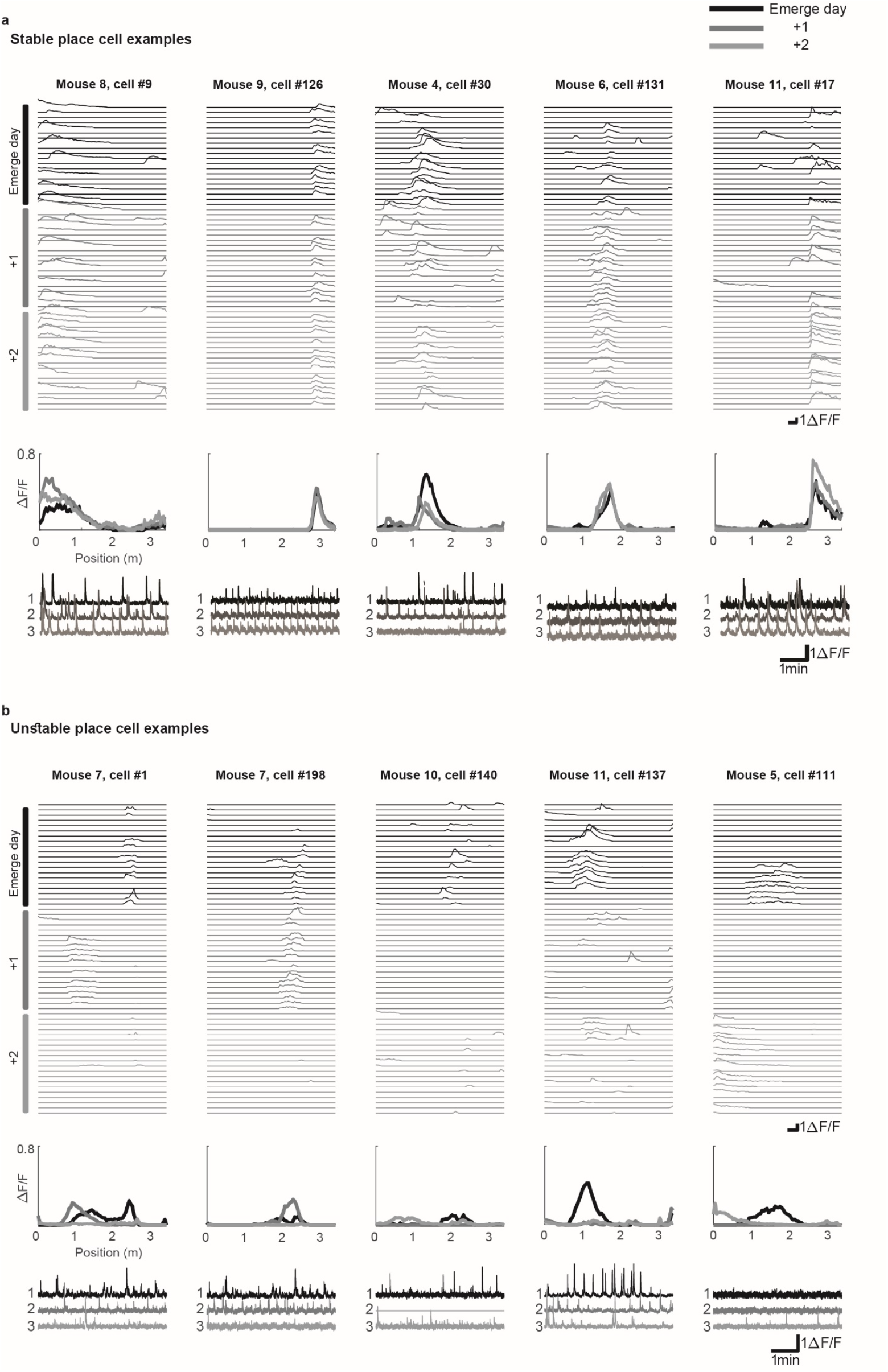
Examples of stable and unstable hippocampal place cells over days. Stable (a) and unstable (b) place cell examples. Top: normalized DF/F trace for the first 10 laps on the emerge day (black) and the two following days (dark and light gray, respectively). Each row is a lap. Middle: mean DF/F vs position (tuning curves) over the three days. Bottom: DF/F time series of each example neuron.

**Extended Data Figure 11:**
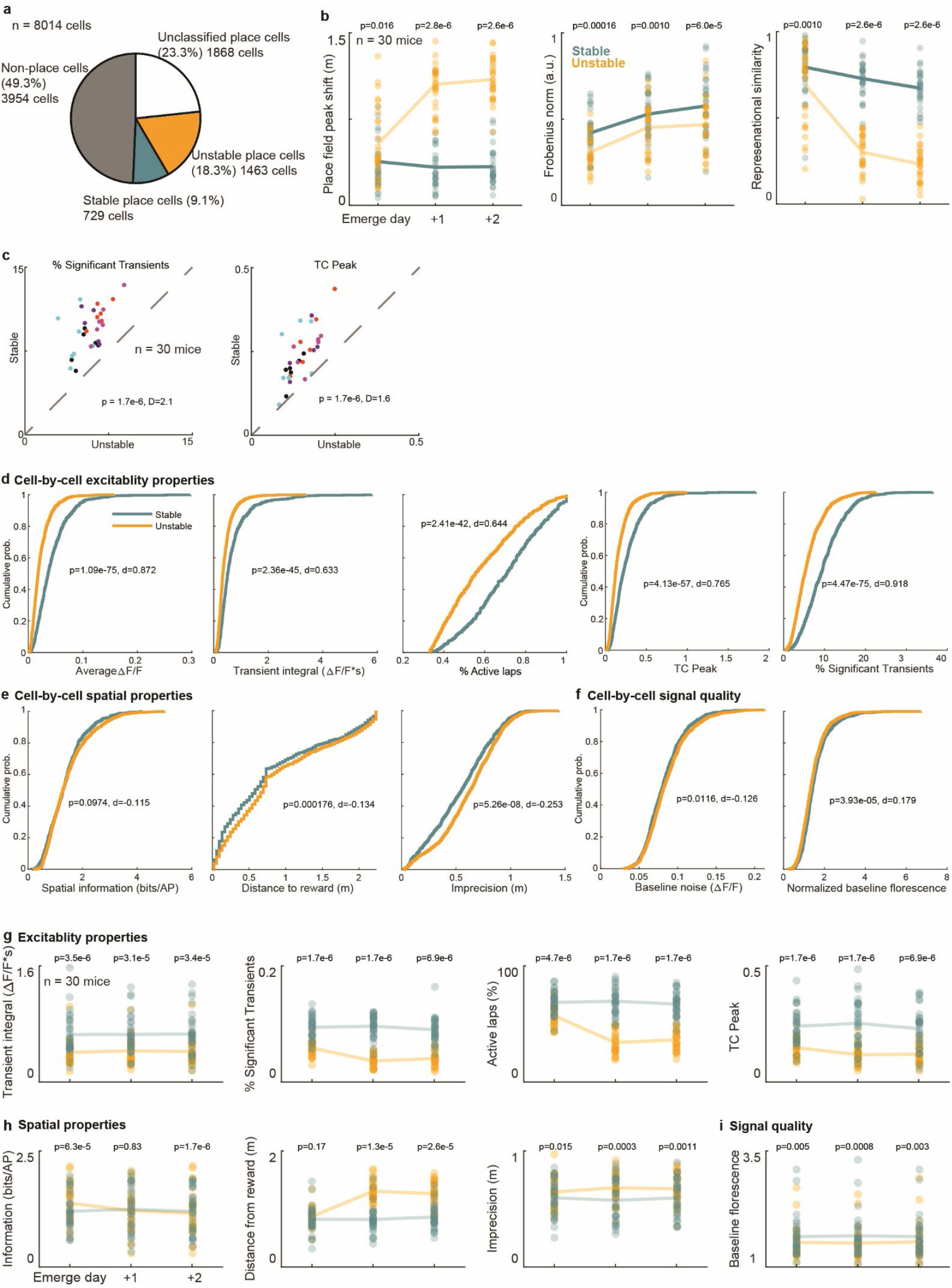
Properties of stable and unstable place cells within and across days. a) Fractions of total pooled neurons by type (n=8014 cells). Unclassified place cells were those that were not categorized into stable or unstable cell categories. b) Additional measures of representational drift for stable and unstable place cells across days. c) Additional excitability properties of stable and unstable neurons. d-f) Cumulative distributions of cell-by-cell excitability properties (d), spatial properties (e), and signal quality (f) for stable and unstable place cells. g-i) Excitability properties (g), spatial properties (h), and signal quality (i) for stable and unstable place cells across days. Wilcoxon rank-sum p values and Cohen’s d are listed for each feature.

**Extended Data Figure 12:**
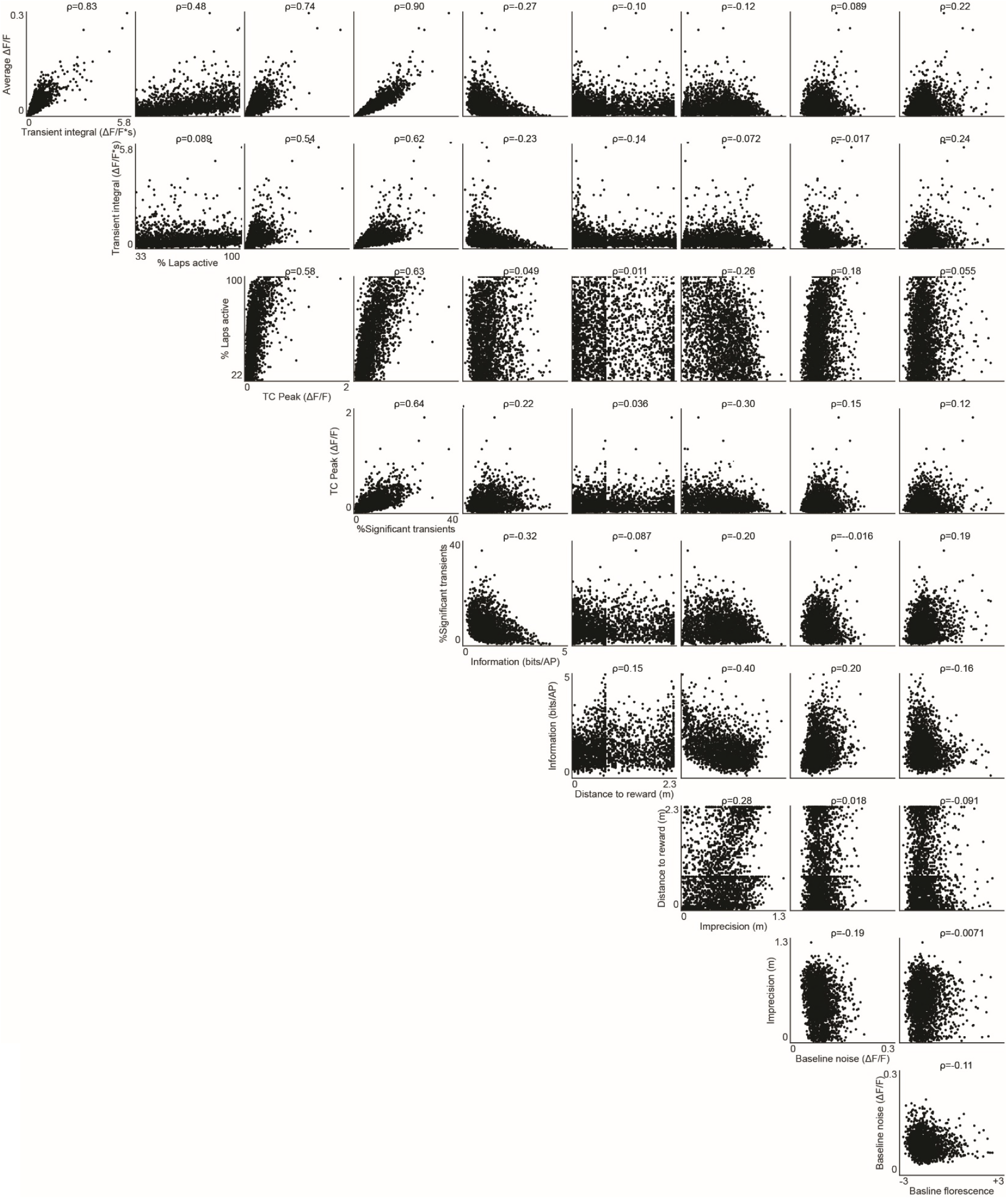
Multicollinearity among various hippocampal neuronal properties of place cells. Pairwise comparisons of each neuronal property with other neuronal properties used in the logistic regression model. Each data point is a single neuron. Pearson’s correlation values for each pair of properties are shown above the panels.

**Extended Data Figure 13:**
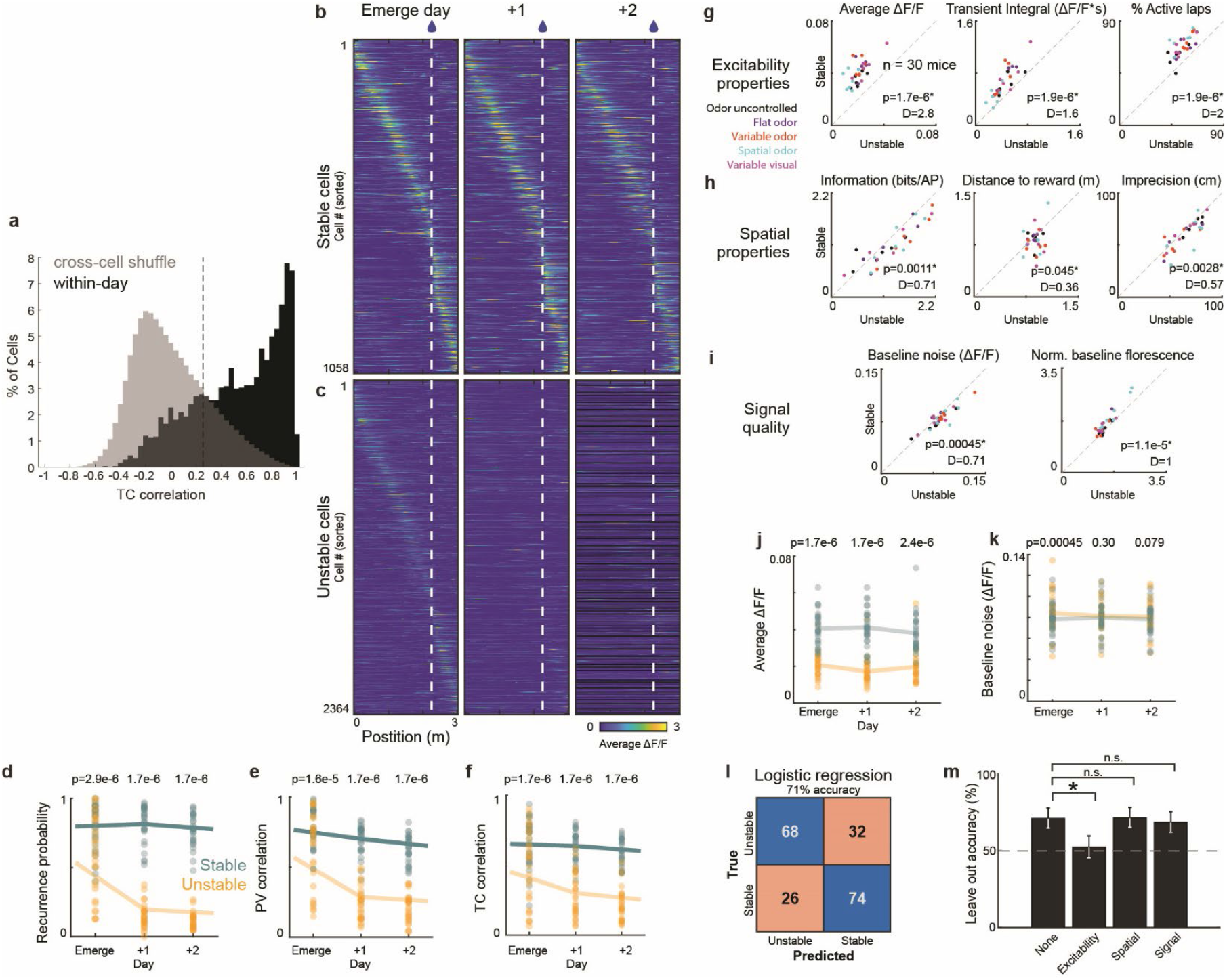
Higher neuronal excitability correlates with and is predictive of more stable representation in hippocampal CA1 across all active cells. a) Tuning curve (TC) correlations for all active cells within day. Odd versus even laps (black) are separated from 10,000 randomly chosen pairs of all active cells (gray) at 0.25 threshold (dashed line). This threshold is used for classification of cells as recurring and therefore, stable (same analysis as the Fig. 4 and Extended Data Figure 2d, but using all active cells instead of only place cells). b-c) Cross-validated heatmaps for all stable and unstable cells from the emerge day and two following days. d-f) Representational drift of stable and unstable cell populations across days (each point represents a mouse, n=30 mice) measured by recurrence probability (d), PV correlation (e), and TC correlation (f) (d-f, sign rank test). g-i) Stable vs. unstable cell excitability properties (g), spatial properties (h), and signal quality (i). Each point is the emerge day average for each mouse (n=30 mice, Wilcoxon signed rank tests and Cohen’s d). J-k) Average dF/F (j) and baseline noise (k) from the emerge day and two following days for stable and unstable cell populations (n=30 mice) (j,k, sign rank test). l) Logistic regression classification of stable vs. unstable cells. The confusion matrix of the fitted regression model predicted whether an active cell becomes a stable or unstable cell from the emerge day properties. m) Accuracy of the logistic regression prediction when each group of properties is left out. Error bars show 95% binomial confidence intervals; likelihood ratio (χ^2^) test.

**Extended Data Table 1:** Detailed statistical values for all ANOVA analysis of the study.

| Figure | Test | Metric | ANOVA type | Effect | F | DF1 | DF2 | p |
| --- | --- | --- | --- | --- | --- | --- | --- | --- |
| Extended Data Fig 2c | Change across days | Total active neurons | 2-way ANOVA | day | 0.18 | 4 | 20 | 0.9455 |
|  |  |  |  | animal | 5.93 | 5 | 20 | 0.0016 |
| Extended Data Fig 2c | Change across days | % Place Cells | 2-way ANOVA | day | 0.33 | 4 | 20 | 0.85 |
|  |  |  |  | animal | 47.43 | 5 | 20 | ~0 |
| Extended Data Fig 2c | Change across days | Place cell information (bits/AP) | 2-way ANOVA | day | 0.72 | 4 | 20 | 0.59 |
|  |  |  |  | animal | 12.86 | 5 | 20 | ~0 |
| Figure 1l | Change across days | Recurrence | 2-way ANOVA | day | 24.46 | 4 | 20 | ~0 |
|  |  |  |  | animal | 2.63 | 5 | 20 | 0.0554 |
| Figure 1m | Change across days | PV Correlation | 2-way ANOVA | day | 22.29 | 4 | 20 | 3.91E-07 |
|  |  |  |  | animal | 10.19 | 5 | 20 | 5.76E-05 |
| Figure 1n | Change across days | TC Correlation | 2-way ANOVA | day | 4.78 | 4 | 20 | 0.0072 |
|  |  |  |  | animal | 2.72 | 5 | 20 | 0.049 |
| Extended Data Fig 2e | Change across days | Peak shift (m) | 2-way ANOVA | day | 5.87 | 4 | 20 | 0.0027 |
|  |  |  |  | animal | 3.99 | 5 | 20 | 0.0113 |
| Extended Data Fig 2e | Change across days | Decoding error | 2-way ANOVA | day | 6.04 | 4 | 20 | 0.0024 |
|  |  |  |  | animal | 13.94 | 5 | 20 | ~0 |
| Extended Data Fig 2e | Change across days | Frobenius norm | 2-way ANOVA | day | 37.22 | 4 | 20 | 5.32E-09 |
|  |  |  |  | animal | 26.13 | 5 | 20 | 3.98E-08 |
| Extended Data Fig 2e | Change across days | Representational similarity | 2-way ANOVA | day | 13.87 | 4 | 20 | 1.42E-05 |
|  |  |  |  | animal | 10.7 | 5 | 20 | 4.09E-05 |
| Extended Data Fig 2e | Change across days | Representational drift index | 2-way ANOVA | day | 21.93 | 4 | 20 | ~0 |
|  |  |  |  | animal | 8.42 | 5 | 20 | 0.0002 |
| Figure 2e | Dissimilar vs similar laps | Within-day speed correlation | 3-way anova | lap set | 21.494 | 2 | 78 | 3.67E-08 |
|  |  |  | Tukeys post-hoc | All laps vs similar laps |  |  |  | 1.17E-07 |
|  |  |  |  | All laps vs dissimilar laps |  |  |  | 5.83E-06 |
|  |  |  |  | Similar laps vs dissimilar laps |  |  |  | 0.61 |
|  |  |  |  | day | 1.78 | 4 | 78 | 0.139 |
|  |  |  |  | animal | 5.02 | 5 | 78 | 0.000483 |
| Figure 2f | Dissimilar vs similar laps | Across day difference | 2-way ANOVA | Lap set | 190.1 | 2 | 10 | 1.11E-08 |
|  |  |  | Tukeys post-hoc | All laps vs similar laps |  |  |  | 2.13E-06 |
|  |  |  |  | All laps vs dissimilar laps |  |  |  | 1.57E-05 |
|  |  |  |  | Similar laps vs dissimilar laps |  |  |  | 6.85E-09 |
|  |  |  |  | Animal | 2.1174 | 5 | 10 | 0.14637 |
| Figure 2i | Dissimilar vs similar laps | Recurrence Prob | 3-way ANOVA | Lap set | 0.076527 | 1 | 49 | 0.78322 |
|  |  |  |  | Animal | 9.5456 | 5 | 49 | 2.10E-06 |
|  |  |  |  | Day | 33.864 | 4 | 49 | 1.49E-13 |
| Figure 2j | Dissimilar vs similar laps | PV Correlation | 3-way ANOVA | Lap set | 1.5607 | 1 | 49 | 0.2175 |
|  |  |  |  | Animal | 22.341 | 5 | 49 | 1.34E-11 |
|  |  |  |  | Day | 24.714 | 4 | 49 | 3.08E-11 |
| Figure 2k | Dissimilar vs similar laps | TC Correlation | 3-way ANOVA | Lap set | 0.096639 | 1 | 49 | 0.75722 |
|  |  |  |  | Animal | 14.242 | 5 | 49 | 1.32E-08 |
|  |  |  |  | Day | 11.991 | 4 | 49 | 7.17E-07 |
| Extended Data Fig 3d | Dissimilar vs similar laps | Peak shift | 3-way ANOVA | Lap set | 0.32178 | 1 | 49 | 0.57313 |
|  |  |  |  | Animal | 5.0573 | 5 | 49 | 8.17E-04 |
|  |  |  |  | Day | 8.2054 | 4 | 49 | 3.79E-05 |
| Extended Data Fig 3d | Dissimilar vs similar laps | Bayesian decoding error | 3-way ANOVA | Lap set | 2.01 | 1 | 49 | 0.16259 |
|  |  |  |  | Animal | 16.267 | 5 | 49 | 1.96E-09 |
|  |  |  |  | Day | 10 | 4 | 49 | 4.49E-06 |
| Extended Data Fig 3d | Dissimilar vs similar laps | Frobenius norm | 3-way ANOVA | Lap set | 0.82628 | 1 | 49 | 0.3678 |
|  |  |  |  | Animal | 56.984 | 5 | 49 | 2.98E-19 |
|  |  |  |  | Day | 56.982 | 4 | 49 | 7.90E-18 |
| Extended Data Fig 3d | Dissimilar vs similar laps | Representational similarity | 3-way ANOVA | Lap set | 0.718 | 1 | 49 | 0.4007 |
|  |  |  |  | Animal | 17.838 | 5 | 49 | 4.89E-10 |
|  |  |  |  | Day | 17.689 | 4 | 49 | 4.80E-09 |
| Extended Data Fig 3d | Dissimilar vs similar laps | Representational drift index | 3-way ANOVA | Lap set | 0.0876 | 1 | 49 | 0.769 |
|  |  |  |  | Animal | 8.976 | 5 | 49 | 4.18E-06 |
|  |  |  |  | Day | 36.24 | 4 | 49 | 4.43E-14 |
| Extended Data Fig 4 | Most vs non-selective pre-lick laps | Recurrence Prob | 3-way ANOVA | Lap set | 5.68 | 1 | 40 | 0.0219 |
|  |  |  |  | Animal | 6.59 | 4 | 40 | 3.58E-04 |
|  |  |  |  | Day | 45.6 | 4 | 40 | 2.18E-14 |
| Extended Data Fig 4 | Most vs non-selective pre-lick laps | PV Correlation | 3-way ANOVA | Lap set | 0.87 | 1 | 40 | 0.35638 |
|  |  |  |  | Animal | 24.17 | 4 | 40 | 3.22E-10 |
|  |  |  |  | Day | 36.51 | 4 | 40 | 7.43E-13 |
| Extended Data Fig 4 | Most vs non-selective pre-lick laps | TC Correlation | 3-way ANOVA | Lap set | 5.48 | 1 | 40 | 0.024 |
|  |  |  |  | Animal | 13.395 | 4 | 40 | 5.16E-07 |
|  |  |  |  | Day | 15.88 | 4 | 40 | 7.29E-08 |
| Extended Data Fig 4 | Most vs non-selective pre-lick laps | Peak shift | 3-way ANOVA | Lap set | 1.22 | 1 | 40 | 0.27 |
|  |  |  |  | Animal | 0.91 | 4 | 40 | 0.467 |
|  |  |  |  | Day | 7.54 | 4 | 40 | 1.26E-04 |
| Extended Data Fig 4 | Most vs non-selective pre-lick laps | Bayesian decoding error | 3-way ANOVA | Lap set | 7.22 | 1 | 31 | 0.01145 |
|  |  |  |  | Animal | 25.16 | 4 | 31 | 1.91E-08 |
|  |  |  |  | Day | 10.55 | 4 | 31 | 1.64E-05 |
| Extended Data Fig 4 | Most vs non-selective pre-lick laps | Frobenius norm | 3-way ANOVA | Lap set | 2.267 | 1 | 40 | 0.13999 |
|  |  |  |  | Animal | 37.66 | 4 | 40 | 4.59E-13 |
|  |  |  |  | Day | 44.58 | 4 | 40 | 3.15E-14 |
| Extended Data Fig 4 | Most vs non-selective pre-lick laps | Representational similarity | 3-way ANOVA | Lap set | 5.02 | 1 | 40 | 0.03059 |
|  |  |  |  | Animal | 13.2 | 4 | 40 | 6.01E-07 |
|  |  |  |  | Day | 23.647 | 4 | 40 | 4.35E-10 |
| Extended Data Fig 4 | Most vs non-selective pre-lick laps | Representational drift index | 3-way ANOVA | Lap set | 2.1 | 1 | 40 | 0.15 |
|  |  |  |  | Animal | 12 | 4 | 40 | 1.68E-06 |
|  |  |  |  | Day | 37.91 | 4 | 40 | 4.15E-13 |
| Figure 3e | Flat vs uncontrolled | Within-day behavioral similarity | Repeated measures ANOVA | Day | 0.888 | 4 | 36 | 0.4809 |
|  |  |  |  | Condition by day interaction | 0.491 | 4 | 36 | 0.7425 |
| Figure 3l | Flat vs uncontrolled | Recurrence probability | Repeated measures ANOVA | Day | 31.98 | 4 | 36 | 2.12E-11 |
|  |  |  |  | Condition by day interaction | 0.48699 | 4 | 36 | 0.745 |
| Figure 3m | Flat vs uncontrolled | PV Correlation | Repeated measures ANOVA | Day | 28.988 | 4 | 36 | 8.15E-11 |
|  |  |  |  | Condition by day interaction | 0.4501 | 4 | 36 | 0.77161 |
| Figure 3n | Flat vs uncontrolled | TC Correlation | Repeated measures ANOVA | Day | 10.299 | 4 | 36 | 1.15E-05 |
|  |  |  |  | Condition by day interaction | 0.90651 | 4 | 36 | 0.4705 |
| Extended Data Fig 5g | Flat vs uncontrolled | Peak shift | Repeated measures ANOVA | Day | 8.8895 | 4 | 36 | 4.24E-05 |
|  |  |  |  | Condition by day interaction | 1.5558 | 4 | 36 | 0.20709 |
| Extended Data Fig 5g | Flat vs uncontrolled | Bayesian decoding error | Repeated measures ANOVA | Day | 7.7655 | 4 | 32 | 1.72E-04 |
|  |  |  |  | Condition by day interaction | 0.41836 | 4 | 32 | 0.79414 |
| Extended Data Fig 5g | Flat vs uncontrolled | Frobenius norm | Repeated measures ANOVA | Day | 44.338 | 4 | 36 | 1.96E-13 |
|  |  |  |  | Condition by day interaction | 0.28112 | 4 | 36 | 0.88827 |
| Extended Data Fig 5g | Flat vs uncontrolled | Representational similarity | Repeated measures ANOVA | Day | 20.266 | 4 | 36 | 8.15E-09 |
|  |  |  |  | Condition by day interaction | 0.87797 | 4 | 36 | 0.48674 |
| Extended Data Fig 5g | Flat vs uncontrolled | Representational drift index | Repeated measures ANOVA | Day | 31.829 | 4 | 36 | 2.27E-11 |
|  |  |  |  | Condition by day interaction | 1.1337 | 4 | 36 | 0.35615 |
| Figure 3e | Flat, uncontrolled and variable odor | Within-day behavioral similarity | Repeated measures ANOVA | Day | 0.98393 | 4 | 56 | 0.42382 |
|  |  |  |  | Condition by day interaction | 0.43186 | 8 | 56 | 0.89696 |
| Figure 3f | Flat, uncontrolled and variable odor | Across days difference score | ANOVA | Condition by day interaction | 0.95197 | 2 | 14 | 0.41 |
| Figure 3l | Flat, uncontrolled and variable odor | Recurrence probability | Repeated measures ANOVA | Day | 78.949 | 4 | 56 | 2.37E-22 |
|  |  |  |  | Condition by day interaction | 1.9641 | 8 | 56 | 0.068151 |
| Figure 3m | Flat, uncontrolled and variable odor | PV Correlation | Repeated measures ANOVA | Day | 62.913 | 4 | 56 | 4.59E-20 |
|  |  |  |  | Condition by day interaction | 2.1477 | 8 | 56 | 0.045969 |
| Figure 3n | Flat, uncontrolled and variable odor | TC Correlation | Repeated measures ANOVA | Day | 24.503 | 4 | 56 | 9.38E-12 |
|  |  |  |  | Condition by day interaction | 1.0544 | 8 | 56 | 0.40777 |
| Extended Data Fig 5g | Flat, uncontrolled and variable odor | Peak shift | Repeated measures ANOVA | Day | 16.13 | 4 | 56 | 7.65E-09 |
|  |  |  |  | Condition by day interaction | 1.3784 | 8 | 56 | 0.22608 |
| Extended Data Fig 5g | Flat, uncontrolled and variable odor | Bayesian decoding error | Repeated measures ANOVA | Day | 21.687 | 4 | 48 | 2.87E-10 |
|  |  |  |  | Condition by day interaction | 2.6314 | 8 | 48 | 0.017804 |
| Extended Data Fig 5g | Flat, uncontrolled and variable odor | Frobenius norm | Repeated measures ANOVA | Day | 89.913 | 4 | 56 | 1.06E-23 |
|  |  |  |  | Condition by day interaction | 1.7329 | 8 | 56 | 0.11087 |
| Extended Data Fig 5g | Flat, uncontrolled and variable odor | Representational similarity | Repeated measures ANOVA | Day | 45.16 | 4 | 56 | 6.68E-17 |
|  |  |  |  | Condition by day interaction | 1.3228 | 8 | 56 | 0.25146 |
| Extended Data Fig 5g | Flat, uncontrolled and variable odor | Representational drift index | Repeated measures ANOVA | Day | 51.606 | 4 | 56 | 3.80E-18 |
|  |  |  |  | Condition by day interaction | 0.86924 | 8 | 56 | 0.54772 |
| Figure 3l | Flat, uncontrolled, variable and spatial odor | Recurrence probability | Repeated measures ANOVA | Day | 98.329 | 4 | 76 | 2.96E-29 |
|  |  |  |  | Condition by day interaction | 1.6072 | 12 | 76 | 0.10743 |
| Figure 3m | Flat, uncontrolled, variable and spatial odor | PV Correlation | Repeated measures ANOVA | Day | 86.386 | 4 | 76 | 1.71E-27 |
|  |  |  |  | Condition by day interaction | 1.8077 | 12 | 76 | 0.061719 |
| Figure 3n | Flat, uncontrolled, variable and spatial odor | TC Correlation | Repeated measures ANOVA | Day | 41.938 | 4 | 76 | 1.59E-18 |
|  |  |  |  | Condition by day interaction | 1.5884 | 12 | 76 | 0.11299 |
| Extended Data Fig 5g | Flat, uncontrolled, variable and spatial odor | Peak shift | Repeated measures ANOVA | Day | 28.177 | 4 | 76 | 2.32E-14 |
|  |  |  |  | Condition by day interaction | 1.3788 | 12 | 76 | 0.19474 |
| Extended Data Fig 5g | Flat, uncontrolled, variable and spatial odor | Bayesian decoding error | Repeated measures ANOVA | Day | 27.876 | 4 | 68 | 1.03E-13 |
|  |  |  |  | Condition by day interaction | 2.2333 | 12 | 68 | 0.019208 |
| Extended Data Fig 5g | Flat, uncontrolled, variable and spatial odor | Frobenius norm | Repeated measures ANOVA | Day | 122.6 | 4 | 76 | 2.41E-32 |
|  |  |  |  | Condition by day interaction | 1.6487 | 12 | 76 | 0.095993 |
| Extended Data Fig 5g | Flat, uncontrolled, variable and spatial odor | Representational similarity | Repeated measures ANOVA | Day | 70.269 | 4 | 76 | 9.04E-25 |
|  |  |  |  | Condition by day interaction | 1.8874 | 12 | 76 | 0.049223 |
| Extended Data Fig 5g | Flat, uncontrolled, variable and spatial odor | Representational drift index | Repeated measures ANOVA | Day | 73.542 | 4 | 76 | 2.32E-25 |
|  |  |  |  | Condition by day interaction | 1.6184 | 12 | 76 | 0.10422 |
| Extended Data Fig 7e | Non-variable visual versus variable visual | Within-day behavioral similarity | Repeated measures ANOVA | Day | 0.61792 | 4 | 44 | 0.65207 |
|  |  |  |  | Condition by day interaction | 0.18962 | 4 | 44 | 0.94257 |
| Extended Data Fig 7h | Non-variable visual versus variable visual | Recurrence probability | Repeated measures ANOVA | Day | 24.794 | 4 | 44 | 8.66E-11 |
|  |  |  |  | Condition by day interaction | 0.50704 | 4 | 44 | 0.73076 |
| Extended Data Fig 7h | Non-variable visual versus variable visual | PV Correlation | Repeated measures ANOVA | Day | 29.034 | 4 | 44 | 7.70E-12 |
|  |  |  |  | Condition by day interaction | 0.37906 | 4 | 44 | 0.82236 |
| Extended Data Fig 7h | Non-variable visual versus variable visual | TC Correlation | Repeated measures ANOVA | Day | 7.739 | 4 | 44 | 8.20E-05 |
|  |  |  |  | Condition by day interaction | 0.48904 | 4 | 44 | 0.74371 |
| Extended Data Fig 7h | Non-variable visual versus variable visual | Peak shift | Repeated measures ANOVA | Day | 6.5863 | 4 | 44 | 3.04E-04 |
|  |  |  |  | Condition by day interaction | 0.73547 | 4 | 44 | 0.57276 |
| Extended Data Fig 7h | Non-variable visual versus variable visual | Bayesian decoding error | Repeated measures ANOVA | Day | 6.9218 | 4 | 40 | 2.48E-04 |
|  |  |  |  | Condition by day interaction | 0.049012 | 4 | 40 | 0.9953 |
| Extended Data Fig 7h | Non-variable visual versus variable visual | Frobenius norm | Repeated measures ANOVA | Day | 63.279 | 4 | 44 | 1.11E-17 |
|  |  |  |  | Condition by day interaction | 0.33545 | 4 | 44 | 0.8526 |
| Extended Data Fig 7h | Non-variable visual versus variable visual | Representational similarity | Repeated measures ANOVA | Day | 15.405 | 4 | 44 | 5.95E-08 |
|  |  |  |  | Condition by day interaction | 0.77383 | 4 | 44 | 0.54814 |
| Extended Data Fig 7h | Non-variable visual versus variable visual | Representational drift index | Repeated measures ANOVA | Day | 45.033 | 4 | 44 | 5.20E-15 |
|  |  |  |  | Experimenter by day interaction | 1.1249 | 4 | 44 | 0.35707 |
| No figure | Experimenter effect | Recurrence probability | Repeated measures ANOVA | Day | 85.24285039 | 4 | 108 | 1.62E-32 |
|  |  |  |  | Experimenter by day interaction | 0.8257427117 | 4 | 108 | 0.5115696947 |
|  | Experimenter effect | PV Correlation | Repeated measures ANOVA | Day | 36.16208707 | 4 | 108 | 3.74E-19 |
|  |  |  |  | Experimenter by day interaction | 1.137927186 | 4 | 108 | 0.3426240207 |
|  | Experimenter effect | TC Correlation | Repeated measures ANOVA | Day | 80.26701289 | 4 | 108 | 1.84E-31 |
|  |  |  |  | Experimenter by day interaction | 1.065738891 | 4 | 108 | 0.3771290116 |
|  | Experimenter effect | Peak shift | Repeated measures ANOVA | Day | 26.25405585 | 4 | 108 | 1.02E-14 |
|  |  |  |  | Experimenter by day interaction | 1.646702138 | 4 | 108 | 0.1688469876 |
|  | Experimenter effect | Bayesian decoding error | Repeated measures ANOVA | Day | 23.38543201 | 4 | 96 | 6.10E-14 |
|  |  |  |  | Experimenter by day interaction | 0.371922104 | 4 | 96 | 0.8282101442 |
|  | Experimenter effect | Frobenius norm | Repeated measures ANOVA | Day | 133.3777017 | 4 | 108 | 7.55E-41 |
|  |  |  |  | Experimenter by day interaction | 1.633523688 | 4 | 108 | 0.1710547551 |
|  | Experimenter effect | Representational similarity | Repeated measures ANOVA | Day | 57.20009643 | 4 | 108 | 8.00E-26 |
|  |  |  |  | Experimenter by day interaction | 0.995244586 | 4 | 108 | 0.4134614623 |
|  | Experimenter effect | Representational drift index | Repeated measures ANOVA | Day | 83.85044921 | 4 | 108 | 3.16E-32 |
|  |  |  |  | Experimenter by day interaction | 1.551837869 | 4 | 108 | 0.1924627742 |

## Methods

### Surgery

All mice procedures were approved by the Northwestern University Institutional Animal Care and Use Committee. Male C57BL/6 mice around ~10 weeks old were anesthetized using 1%–2% isoflurane. To induce the expression of GCaMP8m^24^ in the hippocampus CA1, a volume of 90 nL~125 nl of an adeno-associated virus vector (AAV1.syn.GCaMP8m.WPRE.SV40) with a titer of 2.5 × 10^13 GC/mL was administered via a beveled glass micropipette. A 1 mm diameter craniotomy was performed on the right hippocampus, at a location precisely 1.8 mm lateral and 2.3 mm caudal from bregma. The micropipette containing the virus was carefully inserted to a depth of 1.3 mm beneath the dura surface and the virus was injected. The mice then underwent water restriction, receiving 0.8-1 mL/day, for the duration of the study. Following 2–7 days post-injection, hippocampal window implantation was performed, wherein a stainless-steel cannula featuring a glass coverslip affixed on one side was placed over the hippocampus. Additionally, a titanium head plate and light-blocking ring were securely cemented to the skull as described in previous protocols^21^.

### Multisensory (visual +olfactory) virtual reality apparatus

Mice were head-fixed on a cylindrical treadmill surrounded by a 5-monitor VR display with visual cues constructed in MATLAB using the ViRMEn VR engine^64^. Additionally, the olfactometer system previously described^15,16^ was employed for precise odor control throughout the behavior for the odor-controlled mouse groups. Throughout the training and experiments, we controlled for auditory cues by providing a loud white noise background and were careful to place the nose cone and lick tube in the same location each day to reduce any tactile variability.

In brief, the outlet tubing of the odorant stream led to a 40-mL amber glass vial with a rubber membrane cap (Thermo Fisher) containing 12 mL of odorant solution filled nearly to the top with glass beads. The inlet tubing was submerged in the solution. The outlet tubing in the head space led to a passive mixing chamber where it met a carrier stream of air. The odorant-air mixture flowed through a mass flow controller (MFC) to a final nose chamber covering the mouse’s snout to create an odor micro-environment of 0.07 cm^3. Odorant concentration was controlled by varying the stream flow rate between 0.001-0.100 L/min to achieve relative odorant concentrations between 1%–100%. Odorant solutions included α-pinene (1:37.5 in mineral oil, Sigma-Aldrich) and methyl valerate (1:125 in mineral oil, Sigma-Aldrich) or a mix of both (1:18.74 of α-pinene, 1:62.5 of methyl valerate in mineral oil, Sigma-Aldrich). A waterspout was placed within reach of the mouse’s tongue, and a capacitive circuit (Sparkfun) was used to register licking and was read through a data acquisition card (National Instruments PCI-6229) into MATLAB. All visual cues, MFC flow rates, and water rewards were controlled using the ViRMEn VR engine in MATLAB. Mice received 4 uL water reward two-thirds (2.25 m) of the way along the 3-m virtual track.

### Multisensory VR tasks

#### 1. Uncontrolled odor task

For the “uncontrolled odor” task, no odor and nose chamber were involved in training and experiment days.

#### 2. Variable visual task

In the “variable visual” task, brightness was 100% on Day 1. On Day 2, levels of 86%, 93%, or 100% were randomly selected for each lap. On Day 3, levels of 60%, 67%, or 74% were used at random, and on Day 4, 73%, 80%, or 87%. On Day 5, brightness was fixed at 80%.

#### 3. Flat odor task

For the “flat odor” task, α-pinene odorant with 50% of its maximum flow rate was delivered to mice through the nose chamber during all training and experiment days.

#### 4. Variable odor task

For the “variable odor” task, the odorant type and concentration was different for each day (Fig. 3a-b). On day 1, similar to the “flat odor” mice, α-pinene odorant with 50% of its maximum flow rate was applied. However, for day 2, the odorant concentration was varied on each lap and a randomly chosen 20%, 30%, or 40% of α-pinene odorant was delivered. On day 3, methyl valerate with random 40%, 50%, or 60% concentrations on each lap was delivered. On day 4, α-pinene with random 70%, 80%, or 90% concentrations was delivered. On day 5, a mix of α-pinene and methyl valerate with random 40%, 50%, and 60% concentrations was delivered.

#### 5. Spatial odor task

For the “spatial odor” task, the odor concentration started at 30% at beginning of the track and peaked at the reward location (100%) and tapered off afterward reaching to 76% at end of the track., requiring mice to integrate enriched sensory information and use it to navigate to the reward. This odorant profile was consistent across different laps and days during both training and experiment days.

### Training criterion

To train mice and determine when to start the five-day imaging experiments, we monitored the speed profile of the mice each day. Mice initially learned to slow down and lick before the water reward; however, we relied on the speed profile of the mice for analysis of behavioral stability. There was more behavioral variability when mice were first placed on the track and when they became satiated toward the end of the sessions, so we focused our analysis on the 11th to 30th laps. We constructed a reference lap by taking the mean speed of the mouse in 60 spatial bins across the track. We then used two measures of stability to compare the behavior across days. First, we correlated this reference lap to laps 11-30 on each preceding day. We identified a mouse as trained if the correlation exceeded 0.6 for at least two of the four preceding days. Once these behavioral criteria during training were met, we started the imaging experiments for the subsequent five days. Mice in five experimental tasks experienced their unique task protocols during the training and experiment days.

### Two-photon imaging

A custom movable-objective microscope from Sutter Instruments was utilized for imaging the hippocampal CA1 through a 40x/0.6 NA air objective (LUCPLFLN40x, Olympus) with ScanImage 5.1 software, as previously described^15,16,51^. Time-series movies of 50,000 frames (unidirectional scanning, 31.25 Hz, 512 × 256 pixels movie size) were captured during a ~27-minute behavioral session. A Digidata1440A system (Molecular Devices) via Clampex 10.3 software facilitated data acquisition, recording, and synchronizing factors such as track position, speed, MFC flow rates, licking, reward delivery, and time stamps of each two-photon image frame at 1 kHz.

### Volumetric plane registration of the same imaging field across days

We developed a novel within-session, online pre-experimental, quantitative technique for the longitudinal identification of the same imaging plane across days. On the first imaging day, we acquired 1000 frames of the chosen imaging field with 920 nm and isosbestic 800 nm laser wavelength, motion-correcting these short videos to obtain average images to use as reference templates on subsequent days. In subsequent imaging days, we first identified the putative first-day imaging field qualitatively by visual inspection using previously noted local landmarks. Afterward, we identified the precise imaging field quantitatively using the following methods.

For quantitative identification of the same imaging field in the following days, we captured 15 z-stack slices (200 frames each) with a spacing of 2 um around the putative field. Each z-stack slice was motion-corrected, and the average z-stack slice images were obtained. Each average z-stack slice image was then cross-correlated (using x and y position offsets) with the first day’s template image. For cross correlation, the pixel values of both z-stack slice images and the first day’s template image were standardized to normalized z-scores (center 0, std 1). To determine the best matching XYZ plane, we identified the z-stack slice and the associated x, y offsets that result in the highest peak cross-correlation value. These values therefore reported the positional difference in XYZ coordinates (in microns) between the putative field and the first day’s template image. These values were then used to finely adjust the objective for imaging of the same field as on Day 1. With this technique, we quantitatively and robustly found the same imaging plane across five imaging days with XYZ error of ~2 um.

### ROI detection and fluorescence trace extraction

Registration, ROI selection, and raw fluorescence trace extraction were performed using Suite2p^65^. Once fluorescence traces were extracted, they were converted to Δ*F/F* traces using the 8th percentile method^66^ with a sliding window of 32 seconds (1000 frames). Significant transients were detected using the ratio of positive-to negative-going transients described previously^66^. After the automatic identification of the ROIs, three authors independently evaluated the cellular and subcellular morphologies of every individual cell to validate its correct registration across days. Only the cells that were verified by all three authors were included.

### Place cell spatial properties

Fluorescence tuning maps were made by binning the position across the track into 80 bins and identifying the mean fluorescence where the mouse was moving at least 0.1 cm per second. To test if a cell is a place cell, we computed the spatial information (*I*) in bits per action potential for the fluorescence tuning map^28,67^.

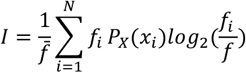

Where 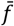 is the mean change in fluorescence, *N* is the number of bins, *f*_*i*_ is the mean fluorescence change in the *i* ^*th*^ spatial bin, and *pX*(*x*_*i*_) is the probability that the animal is in the *i i* th spatial bin.

To build a null distribution of information, we circularly shuffled the fluorescence trace with a minimum shift of 15 seconds and recalculated the tuning map 1000 times. A cell was a significant place cell if it had higher information than 99% of these shuffle epochs, had an information of at least 0.5 bits per action potential, and had at least one significant transient during running on at least ⅓ of the laps.

Place cell “imprecision” was measured as the standard deviation of the peak DF/F location on active laps. Place field “distance to reward” was defined as the mean of distances of place field peak to the reward location on the virtual track.

We defined “stable” cells as place cells that recurred on 4-5 consecutive days and “unstable” cells as place cells that recurred for two or fewer consecutive days and did not recur on the first or the last day.

### Quantification of representational drift

To ensure that mice were engaged with the task while we measured neural activity, we only used laps where mice slowed before the reward. To do this, we compared the speed in two different 30 cm zones, one just before the reward and the other 70-100 cm from the start of the track, where the mice are typically running full speed. We restricted our analyses to laps where the mouse was running in the reward zone at <50% the speed from the zone earlier on the track. This resulted in 64.1±2.83 laps used per session out of 77.5±2.46 total laps per session (82.3±2.25% of laps included per session).

We used eight quantitative measures of representational drift:

1. Recurrence probability (either place cells only or all active cells) A cell was considered recurrent between two days if it was active on ≥1/3 of laps on both days and had a tuning curve (TC) correlation >0.4 for place cells. This threshold was set via logistic regression, comparing within-day TC correlations of place cells to 1,000 randomly paired cells (Extended Data Figure 2d). For stable or unstable place cell classification across days (Fig. 4), this measure was used. For the analysis comparing stable versus unstable cells within the active cell population (all cells active for at least 1/3 of laps on both days; Extended Data Figure 13), the TC threshold was determined to be 0.25 using logistic regression following the same procedure (Extended Data Figure 13a).
2. Population vector (PV) correlation (all cells) This measure quantifies how similarly a neuronal population is activated by taking the median PV correlation across the track.
3. Tuning curve (TC) correlation (place cells only) This measure correlates each neuron’s mean ΔF/F vs. position (i.e., tuning curves, TC) across days. Only cells active on all five days and with non-zero variance were included. We further restricted this analysis to cells that met the place cell criterion on Day 1.
4. Place field peak shift (place cells only) For cells active on all five days, the shift in the peak of their tuning curves across days was calculated.
5. Bayesian decoding error (all cells) Following a previously described method^68^, Bayesian decoding was performed on the likelihood that a significant transient occurred in a 0.1 second bin (*Δt*). The conditional likelihood that an animal is in position *x*_*i*_ given the number of active frames during a time window (*n*) is:

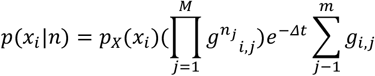

Where *g*_*i, j*_ is the average rate of significant frames by the *j*th neuron in the *i* th spatial bin, *n*_*j*_ is the number of significant frames observed during the time window in neuron *j*, and *M* is the total number of neurons. The decoded position was selected as the one with the maximum conditional likelihood. Because the number of neurons significantly impacts the quality of decoding, the position was chosen as the average position decoded through 100 random samples of 150 neurons: if a mouse had less than 150 neurons, it was excluded from this analysis. The error was determined as the mean error across the session.
6. Frobenius norm (all cells) The Frobenius norm of the peak-normalized spatial maps was calculated, then normalized by the square root of the number of cells.
7. Representational similarity (all cells) Calculated as the Pearson’s correlation coefficient between linearized spatial maps, as reported previously^20^.
8. Representational drift index (all cells) The representational drift index (RDI)^20^ was computed as:

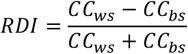

Where *C*_*ws*_ and *C*_*bs*_ are the average correlation coefficient of the linearized spatial maps for all laps within a session and across sessions respectively. For the within-day comparisons, odd and even laps were used.

### Separation of similar and dissimilar running behaviors

#### “Similar” running set

To find laps with similar speed profiles across days, we first created a speed vector for each lap that passed the slowdown criterion using 60 spatial bins (1 bin = 5 cm). To find the laps that were maximally similar across days, we used a particle-swarm and clustering technique. The particle swarm was initialized with all the laps that passed the slowdown criterion. For a given particle, we found the 20 laps on each day that minimized the Euclidean distance of the speed vector to that particle. We then computed the mean Euclidean distance of these laps to the particle. The swarm was optimized to minimize this distance, identifying laps that were similar to each other across the five days.

#### “Dissimilar” running set

To identify laps that were dissimilar across days we maximized the amount of variance explained by the day using the particle swarm optimizer. Reference laps were chosen for all five days, and the 20 laps closest by Euclidean distance within day were identified. We then ran a 3-way ANOVA across day, position, and their interaction, treating each spatial bin as an independent observation. The fraction of variance explained by the day (*R*_*day*_) was taken as:

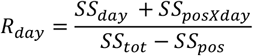

Where *S*_*day*_ is the variance explained by the day, *SS*_*posXday*_ is the variance explained by the position-by-day interaction, *SS*_*tot*_ is the total variance, and *SS*_*pos*_ is the variance explained by position. The particle swarm optimizer was used to maximize this fraction. The optimizer was seeded with 250 pseudo-randomly selected laps from each day.

We then validated that the laps identified by this swarm analysis were indeed similar/dissimilar by using two ways to quantify the amount of behavioral variability. First, to examine how stable the selected behavioral laps were within a day, we found the mean correlation of the position binned speed vectors on selected laps to the average on each day. Second, we used PCA to reduce the speed vector space to three dimensions, which contained 63.9±3.5% of the variance. We then took the normalized distance between the center of the laps chosen each day.

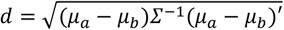

Where *µ*_*a*_, *µ*_*b*_ are the centers of the laps chosen on days *a* and *b* and *Σ* is the covariance matrix for all laps. The difference score was taken as the mean pairwise normalized distance for all combinations of days. To generate a shuffle for this distance, 1000 random pairs of laps were chosen, and the normalized pairwise distance was calculated.

### Most selective and non-selective pre-lick behaviors

We quantified the degree of pre-licking per lap using a pre-lick index^36^, calculated as the difference between the average lick rate in the 30 cm before the reward site and the average lick rate on the rest of the track, divided by their sum. For the most selective pre-lick set, we used the 20 laps with the highest scores above 0. For the non-selective set, we used up to 20 laps with scores below 0. Mice were included only if they had at least 10 non-selective pre-lick laps on Day 1 (to allow even-odd comparisons) and 5 on subsequent days.

### Internal tuning curves

To compare internal structure to spatial tuning we created internal tuning curves. First, Fc3 traces were binarized and then were mapped onto the first 4 dimensions using the non-linear dimensionality reduction algorithm Laplacian Eigenmaps^99^ as previously described^69^. These data were clustered into 10 states using k-means clustering. No sub-clustering was performed. The order of states was defined using transition matrix algorithm described by Rubin *et al*. ^69^, with the exception that the “first” state was determined as the state with median timestamp closets to the start of trial, and that the direction of movement through the states was determined as the direction which maximized the correlation between state and time since start of trial. Significant state cells were identified as having significant state information (see Place Cell Spatial Properties) although no minimum threshold of information was used. Significant cells were sorted according to their state-peak during epochs that corresponded to odd trials, and tuning curve maps were made for the even trials. These were visually compared with spatial maps made from the full running epochs with the same cell selection and sorting criterion.

### Imaging signal quality measures

Baseline noise was estimated using the standard deviation of the *ΔF/F* trace outside of significant transients. Baseline fluorescence was calculated as the average extracted fluorescence in the first 10 seconds of the recording outside of significant transients, normalized by the median intensity of the mean imaging field.

### Logistic regression

We computed several potential predictors of place cell stability (Table 2). Because some of the predictors followed lognormal distributions^69^, they were first transformed into log space. Moreover, before training the model all data was standardized. To identify the best predictors of place cell stability, we used an elastic net logistic regression approach^70^. First, we randomly chose 100 stable and 100 unstable cells to be held out of training. We used class weights to counterbalance the number of observations in each class in the training data set. We used a grid search of 200 λ values between 10^−6^ and 10^6^ on a logistic scale and 100 α values between 0 and 1, selecting the values with the lowest deviance under 5-fold cross-validation in the training set. To test model performance, we used predicted values above 0.5 as stable and measured the accuracy of the predictions on the held-out data in a balanced test set (number of correct predictions/number of held-out data points).

**Table 2:** Predictors and transforms applied before logistic regression.

| Predictor | Log transform | Predictor | Log transform |
| --- | --- | --- | --- |
| Average $\Delta F/F$ | Yes | Information (bits/AP) | No |
| Transient integral ($\Delta F/F \cdot s$) | Yes | Distance from reward (m) | No |
| % Laps Active | No | Imprecision (m) | No |
| Tuning Curve Peak ($\Delta F/F$) | Yes | Baseline noise ($\Delta F/F$) | Yes |
| Time active (%) | No | Baseline fluorescence ($\sigma$) | No |

### Quantification and statistical analysis

The data were analyzed using custom MATLAB codes. Sample sizes weren’t predetermined statistically, and the experimenter was aware of the experimental tasks. Inclusion criteria were based on the behavioral metrics detailed in the Methods section. Non-parametric and parametric statistical tests were used as indicated in the main text and methods. We have reported statistical parameters, such as the precise value and interpretation of ‘n’, precision indicators (represented as mean or median ± SD or SEM), and the establishment of statistical significance, in both the main text and accompanying figure legends. The significance threshold was set at p < 0.05.

## Data availability/Code availability

Matlab code and datasets will be posted to online sharing servers. Additionally, all Matlab codes used in this project can be obtained upon request. For additional information, or requests pertaining to resources and reagents, please reach out to the lead contact, Daniel Dombeck.

## Ethics declarations

The authors declare no competing interests.

## Notes

### Competing Interest Statement

The authors have declared no competing interest.

### Funding

- National Institutes of Health: T32AG020506, 1F32NS116023, R01MH101297

